# Modeling Pyramidal Neurons Using Bidomain BEM and Hierarchical Matrix Approximation

**DOI:** 10.1101/2025.02.21.639592

**Authors:** Nahian I. Hasan, Vanine Sabino, Amanda Jeanne Walenciak, Yang Liu, Luis J. Gomez

## Abstract

Electromagnetic brain stimulation uses electrodes or coils to induce electric fields (E-fields) in the brain and affect its activity. Our understanding of the precise effects of the device-induced E-fields on neural activity is limited. In this paper, we present a novel bidomain boundary integral equation-based method that enables the modeling of fully coupled E-fields from both neurons and stimulation devices. This boundary element approach is accelerated using fast direct solvers to allow for the analysis of realistic scenarios. We present examples, indicating the ability of our solver to analyze rat L2/3 pyramidal neurons derived from the Blue Brain Project. A comprehensive analysis shows that this method can be easily extended to model a group of neurons that were previously computationally intractable.

## I. Introduction

In electromagnetic brain stimulation, coils or electrodes driven by low-frequency current pulses induce electric (E) fields in the brain. Correspondingly, these E-fields affect brain activity. To determine the effects of E-fields on the brain, an important first step is to characterize their effects on single neurons and small networks of neurons. Here, we introduce a scalable direct solver accelerated boundary element bidomain approach to determine the effects of the E-field on neurons and small networks of neurons.

Standard approaches to analyze neural activity induced by E-fields use computational E-field dosimetry results as a forcing function (e.g., activation functions [1] or quasi-potentials [2]) that drives a filamentary cable model of a neuron. The underlying assumption is that a cell and its surrounding cells do not affect the E-field observed in the extracellular space, as such, the device E-field is introduced as an extracellular source driving the circuit [1]–[6]. This source is then incorporated into cable models of neuron cells as distributed current or voltage injections. However, this assumption is known to be incorrect. Local extracellular E fields around neurons are significantly modified by transmembrane ion currents [7]. Additionally, it was recently found that surrounding cells distort the device-induced E-fields, thereby changing the amount of E-field required to activate a neuron(i.e., neuron activation thresholds) [8]–[10].

Recently, [9] proposed using boundary element fast multipole method (BEM-FMM) [11], [12] to solve for the E-fields in the presence of cells and use these corrected E-fields as an activating function. Their approach offers an improvement over standard methods, however, it neglects changes in cell membrane electrical properties resulting from ionic channels opening and closing. Another approach, uses finite element method (FEM) to solve the full coupling between the cell and its surroundings [1], [13]–[15]. While these FEM-based methods capture the full coupling between the cells and surrounding tissue, they require volumetric meshes that bridge individual neurons (micrometers) to whole-brain (centimeters) scales. The generation of multi-scale meshes can make these bidomain FEM methods impractical or intractable for more complex simulations. As a way to avoid volumetric meshes, we introduced a bidomain boundary element method (BEM) for determining the neuron’s response to an impinging E-field [10]. This approach avoids the multi-scale limitations of FEM while retaining full coupling. Unfortunately, the execution of this solver requires the solution of a dense matrix system of equations for each time-step, and a typical simulation has thousands of time-steps. As such, the bidomain BEM requires an intractable level of computational resources for analysis of realistic scenarios.

In this paper, we extend the bidomain BEM method by accelerating it using a direct solver. The dense BEM matrix is highly compressible, and we utilize the hierarchical offdiagonal low-rank (HODLR) approximation technique [16] to represent the interactions between different sub-groups of charges on the membrane surface with a minimal number of basis functions. This allows a precomputation of the inverse of the HODLR compressed BEM matrix for a large system and becomes a simple matrix-vector product routine during the computation and update of the membrane voltage. Our approach is the first scalable tool that can model E-field effects on neuron cells without including severe simplifications. We provide results demonstrating the scalability of our technique along with results from layer (L) 2/3 pyramidal neurons and benchmark the results for a large group of neurons.

## II. Methods

### A. Adjoint BEM Method

We consider a cell inside an inhomogeneous conductive medium stimulated by electrodes or coils (Fig.[1]). The E-field generated by the electrodes and coils is assumed to have a low-frequency spectrum (≤ 100 kHz), allowing for the validity of quasi-stationary assumptions [17]. The electrodes are driven by a known current 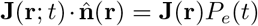 that is only non-zero on the electrodes (i.e., cathodes and anodes). Here, *P*_*e*_(*t*) is the driving current pulse waveform, and 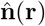 is the outward-pointing unit normal vector. The coil is driven by a known current **J**_**coil**_(**r**; *t*) = **J**(**r**)*P*_*c*_(*t*), where *P*_*c*_(*t*) is its driving pulse waveform. The coil generates an E-field 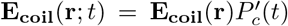 in empty spaces, where 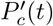 is the time derivative of the current pulse to the coil. This results in a charge distribution *ρ*(**r**; *t*) accumulated on boundaries between media with distinct conductivities. This accumulated charge results in a secondary E-field **E**_*ρ*_(**r**; *t*). In addition, the cell has a thin membrane that mediates the exchange of ions between its intracellular and extracellular spaces. This membrane has a voltage drop of 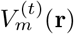 across it that results in an E field **E**_**m**_(**r**; *t*). The adjoint double layer is derived by imposing continuity of the normal component of the conduction current at boundaries between media with distinct conductivities and the cell membrane as

**Fig. 1:**
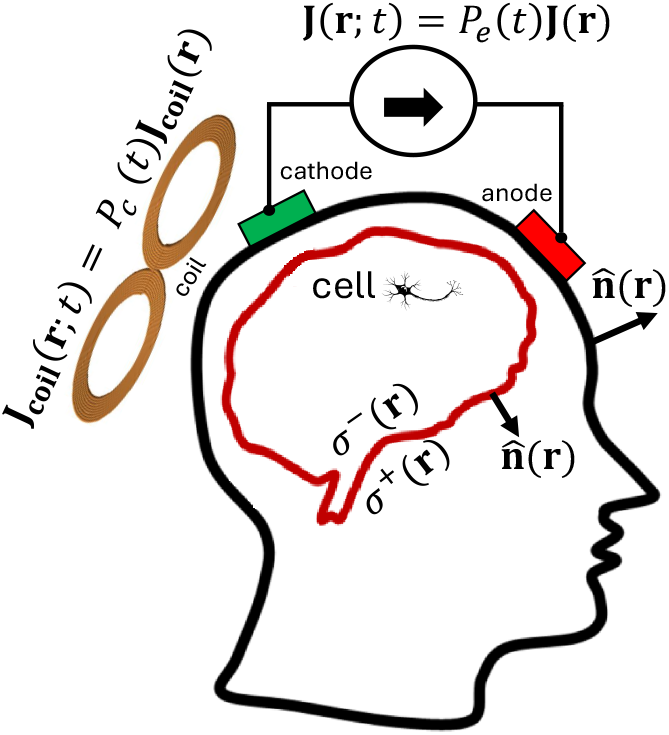
A cell inside the inhomogeneous conductive medium (head) being stimulated by electrodes (cathode and anode) and a coil.

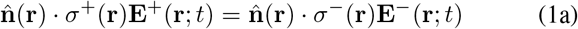

for {**r** ∉ *electrodes*} and

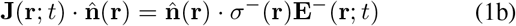

for {**r** ∈ *electrodes*}.

Here, *σ*^+*/*−^(**r**) and **E**^+*/*−^(**r**; *t*) are the conductivities and E-fields just outside/inside of the boundary, respectively. Equations (1a) and (1b) are writen more compactly as

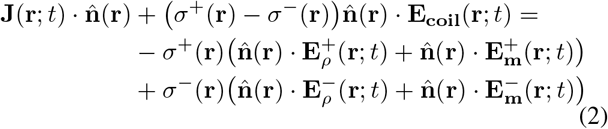

by setting *σ*^+^(**r**) = 0 wherever **r** ∈ *electrodes*. The three components of the total E-field inside the head are

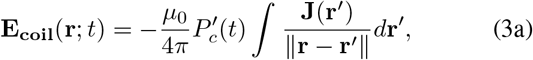

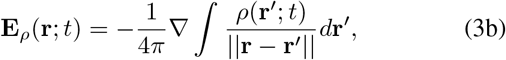

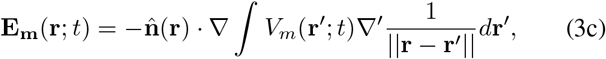

where *µ*_0_ is the permeability in free space. A more detailed and generalized mathematical derivation of the bidomain BEM can be found in [10].

Both the membrane voltage and charge density are unknowns. However, the membrane voltage is related to the membrane current *I*_*m*_(**r**; *t*), which again is proportional to the charges as

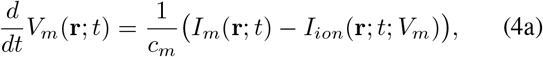

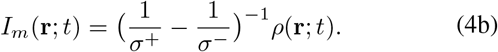

Here *I*_*ion*_(**r**; *t*; *V*_*m*_) is the total flow of current due to the exchange of charged ions between the intracellular and extracellular space, *c*_*m*_ is the membrane capacitance. The ionic currents can be determined from the membrane voltage and equivalent circuit models of the cell membrane. To find the ionic currents from the membrane voltage, we utilize distinct circuit models for the soma, axon, and apical and basal dendrites given in Section II-F.

To determine the membrane voltage, current, and charges, we approximate the boundaries of the cell and surrounding media as a triangle mesh consisting of *N*_*T*_ triangles and *N*_*V*_ nodes. All quantities are approximated using piece-wise linear nodal elements on the triangle mesh as 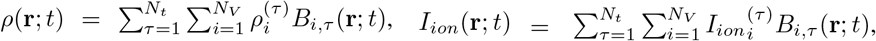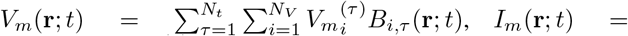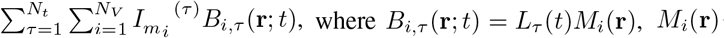 is the nodal element corresponding to the *i*^th^ node of the mesh and is a hat basis function centered at *t* = *τ* Δ*t* and with support (*τ* − 1)Δ*t < t ≤ τ* Δ*t* and *N*_*t*_ is the number of time-steps. It is assumed that we know the state of all quantities at the initial time *t* = 0.

We use (4a) and (2) to determine all quantities at time (*τ* + 1)Δ*t* from all quantities at time *τ* Δ*t*. A semi-implicit approximation of (4a) is made and evaluated at each node of the mesh as

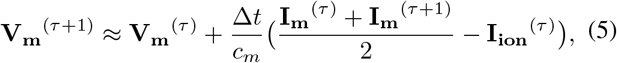

where 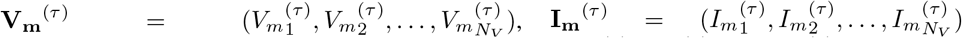, and 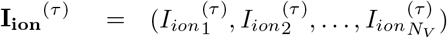. The right-hand side of (5) and (4b) are used to cast (1) as an integral equation with 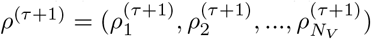 as unknown. A standard Galerkin approach is applied to the resulting equation, and this results in the following linear system of equations

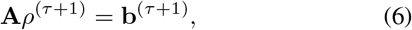

where

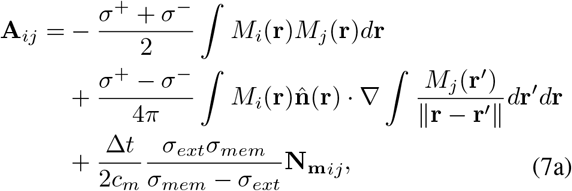

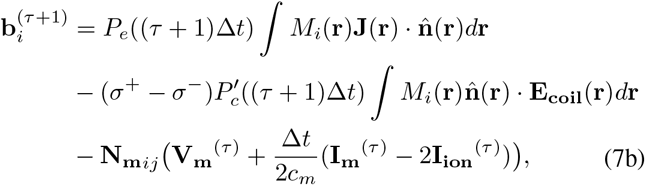

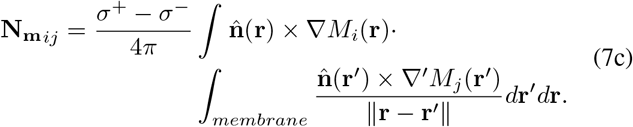

Here *σ*_*mem*_ is the cell membrane conductance, *σ*_*ext*_ is the conductivity of the region surrounding the cell. The entries of the matrix **A** that correspond to 3^rd^-order neighbors are computed using analytical [10], [18], [19] formulas for the inner integral and a 8^th^-order accurate quadrature rule for the outer one. In other words, if the column and row indexes correspond to nodal elements with corresponding nodes connected on the mesh by a path of three edges or fewer, they are computed to high accuracy. All other matrix entries are approximated using single-point triangle centroid quadratures. The system of equations is solved to determine *ρ*^(*τ*+1)^. Once *ρ*^(*τ*+1)^ is known, the membrane current coefficients **I**_**m**_^(*τ*+1)^ are determined using (4b), and **V**_**m**_^(*τ*+1)^ using (5). Finally, **I**_**ion**_^(*τ*+1)^ is computed from **V**_**m**_^(*τ*+1)^ using (12) whose evaluation is described in section II-F.

The storage of **A** requires order 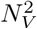 memory. Correspondingly, this limits the complexity of scenarios that can be studied with the standard Bidomain BEM. Here we compressed the matrix **A** and stored it using a hierarchical off-diagonal low-rank (HODLR) format requiring only order *N*_*V*_ *log*(*N*_*V*_) memory. Furthermore, we describe an approach to compute its LU factors in the HODLR format, enabling its fast-direct solution.

### B. HODLR compression of A

To lower the memory complexity of storing the matrix **A**, its LU factors are stored in a HODLR format. The HODLR matrix **H** ≈ **A** is stored to a prescribed accuracy ||**A** − **H**|| *< ϵ*_*H*_ || **A** ||, where *ϵ*_*H*_ is a user specified error level (here chosen as 10^−7^) and || · || is the Frobenius matrix norm. A two-level HODLR matrix (**H**) is shown in Fig. [2]. In Fig. [2], compression is achieved by only storing full-diagonal sub-matrices (shown in red) and compressing the others. Several methods can be used to compress the off-diagonal submatrices; here we used adaptive cross-approximation (ACA) [20]–[22], and its details are given in the next section. Next, the hierarchical structure of the HODLR matrix is described in more detail.

**Fig. 2:**
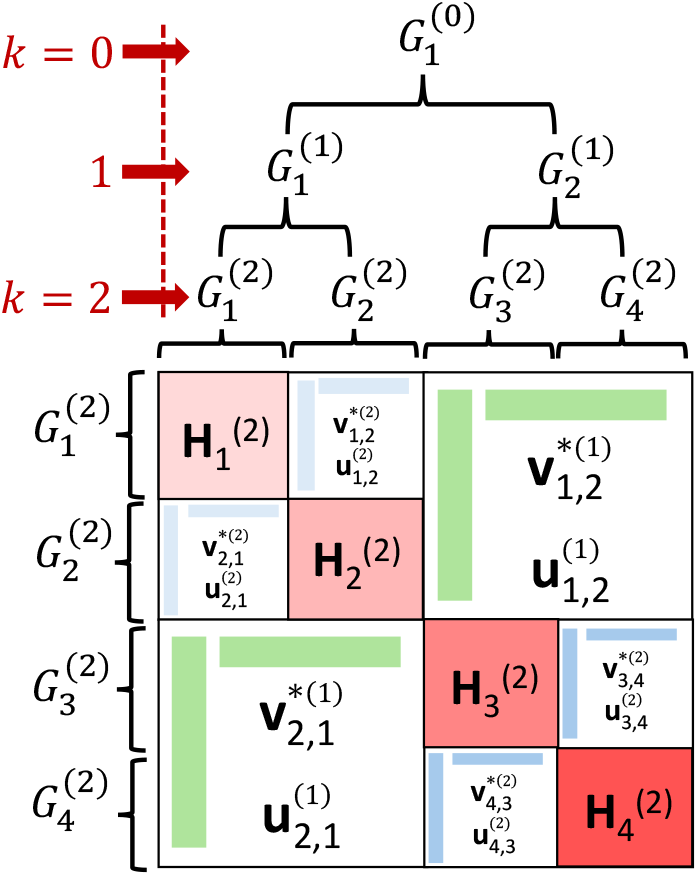
HODLR matrix for 2 levels of hierarchical partitioning. The *k*^th^ level has 2^*k*^ groups of nodes. The off-diagonal admissible blocks are computed using a low-rank approximation technique. At the last level (*k* = 2), the diagonal inadmissible dense blocks 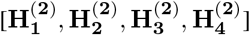 are shown in solid red colors.

The construction begins by generating an *L*-level binary tree partitioning of the row/column indices based on their corresponding node coordinates. At the first level of the tree, nodes are divided into two groups by choosing a bisecting plane and assigning nodes that are on the same side of it to the same group. The bisecting plane is chosen to cut the axis-aligned minimum bounding box in half along its longest dimension, also known as the KD-tree partitioning. At the second level, each of the two groups from the first level is divided using the same method, resulting in a partition into four groups. This process continues recursively for *L* levels, where each level doubles the number of groups. At the final level, the mesh is divided into 2^*L*^ groups, ensuring a hierarchical partition structure. The groups are arranged hierarchically to maintain inheritance relationships. Specifically, each group at level *k* consists of two child groups of level *k* + 1. Furthermore, the *i*^th^ group from level *k* is formed by merging the groups 2*i* − 1 and 2*i* from level *k* + 1 and is referred to as its parent. (Note: Without loss of generality, we assume that the nodes are numbered in an ordered way with an increasing node index corresponding to an increasing group index. In other words, at the *L*^th^ partitioning level, the indexes of the nodes in the *i*^th^ group are 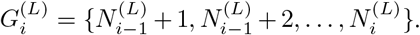. We define 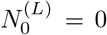 and 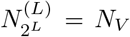 and 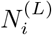 to be the index of the last element of the *i*^th^ group at level *L*. As a result, the nodes in the *i*^th^ group at level *k* have indexes 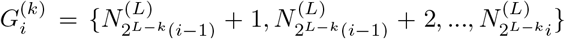. This numbering scheme ensures that each group at a higher level is a direct aggregation of its corresponding child groups, preserving a structured hierarchy throughout the partitioning process.

The HODLR submatrices are selected and formatted according to simple rules. At each level, matrices corresponding to row and column indices of two groups sharing the same parent are stored using ACA. In other words, 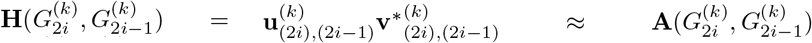 and 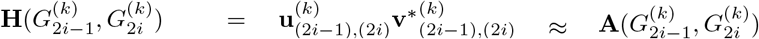, where *k* ∈ {1, 2, …, *L*}, *i* ∈ {1, 2, …, 2^*k*−1^}. At the lowest level, submatrices where the row and column indices belong to the same group are stored in a dense format. In other words, 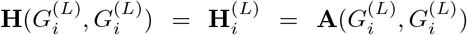. For example, in Fig.[2], at level 1, submatrices 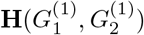 and 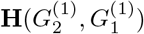 are compressed and its factors are shown in green. At level 2, compression applies to submatrices 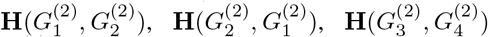 and 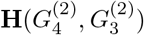 as shown in blue. Additionally, at the second level, all diagonal submatrices are stored in full uncompressed format as shown in red. For interested readers, a 3-level HODLR matrix and its LU decomposition are provided in the supplementary file.

## C. Low-rank approximation of off-diagonal blocks

The HODLR compression relies on special properties of the **A**. In particular, it is well known that its off-diagonal blocks have an *ϵ*-rank that scales as *log*(*ϵ*) under the quasi-stationary assumption [23]. The *ϵ*-rank of a matrix is the smallest number of singular values required to approximate it to *ϵ* accuracy (i.e., *r* is the *ϵ*-rank if *r* is the smallest integer such that 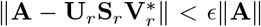, where **U**_*r*_, **S**_*r*_, **V**_*r*_ are a singular value decomposition consisting of the the first *r* singular vectors and values. A similar decomposition can be found from linearly independent columns and rows of **A**. For example, the interpolatory decomposition or CUR 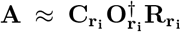, where 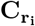 are *r*_*i*_ linearly independent columns, 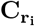 are *r*_*i*_ linearly independent rows, 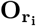 is a matrix composed of the overlap matrix between the *r*_*i*_ rows and columns,and denotes pseudo inverse. Whenever the rows and columns are chosen to maximize the determinant of 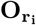, this decomposition has been shown to produce practically nearly optimal compression. Furthermore, unlike the SVD, its construction in principle does not require the evaluation of the whole matrix. Here we employ the ACA algorithm to compress off-diagonal blocks of the HODLR matrix without needing to compute the full off-diagonal blocks. The ACA iteratively generates an interpolatory decomposition of a matrix **A**, increasing the rank of the approximation by 1 at each iteration by appending a column and row, which will maximize the correction based on the current approximation. The ACA [24] was introduced to compress BEM matrices and has been very successful [21], [25], [26]. In particular, the ACA factorizes the rows and column of the matrix to write an equivalent form of the interpolatory decomposition involving only two factors 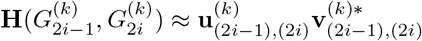.

### D. Computing the LU of the HODLR matrix

The system of equations described in (6) must be solved at each time step. Here we describe the method we use to decompose **H** into lower-triangular **L** and upper-triangular **U** matrices **H** = **LU** in the HODLR format and the inverse **H**^−1^ = **U**^−1^**L**^−1^. The charge distribution at each time-step can be determined using the inverse LU factors as *ρ*^(*τ*+1)^ = **U**^−1^**L**^−1^**b**^(*τ*+1)^. The construction of the **U**^−1^ and **L**^−1^ only requires order 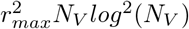 computations, where *r*_*max*_ is the maximum compression rank; thereby reducing the computational run-time. The algorithm for computing the LU decomposition of an HODLR matrix is mentioned in [27]. Specifically, at level *L*, the LU factorization 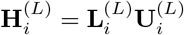 of each of the diagonal dense blocks are computed using LAPACK routines. The LU factorization for parent groups is computed using the following formula

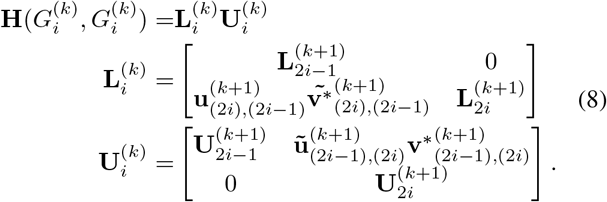

where 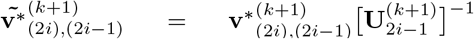 and 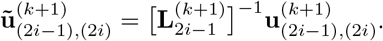. (Note: The algorithm as explained has a computational cost of order 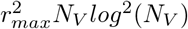 [16], [27]. Fig. [3] illustrates the LU factorization of the two-level matrix of Fig. [2]. At level 2, the LU factors are computed for each of the red blocks (Fig.[3A]). At the first level, the LU factorization of each group is computed by concatenating its children group LUs and generating new compressed off-diagonal blocks (Fig. [3B]). The final structure of the LU factors is shown at the bottom of the figure.

**Fig. 3:**
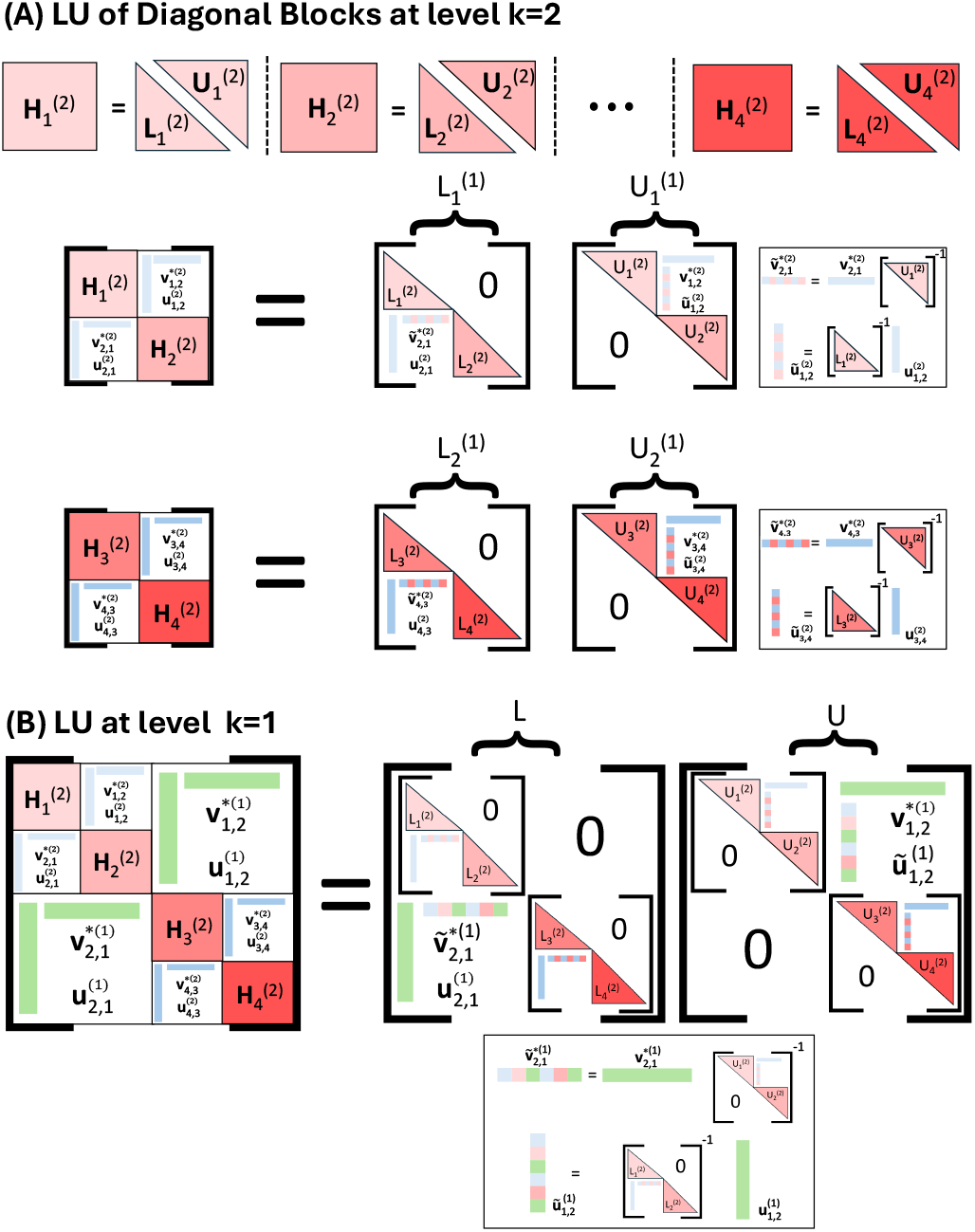
LU decomposition of the HODLR matrix. (A) Diagonal inadmissible blocks at level *k* = 2 are factorized using direct LU decomposition. (B) LU decomposition at level *k* = 1.

To compute the inverse of each of the LU factors at the lowest level, the inverse of the 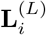 and 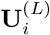 factors is computed using standard methods. Then, the other levels are computed as

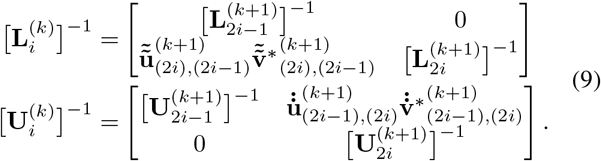

where 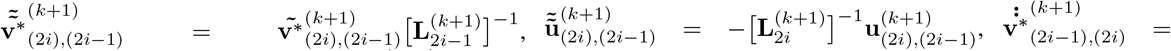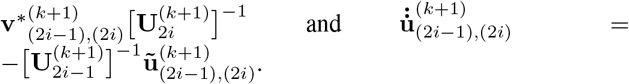.

A summary of the bidomain BEM algorithm is shown in Fig.[4], which depicts the algorithmic flow chart including HODLR compression and updating the gating variables.

**Fig. 4:**
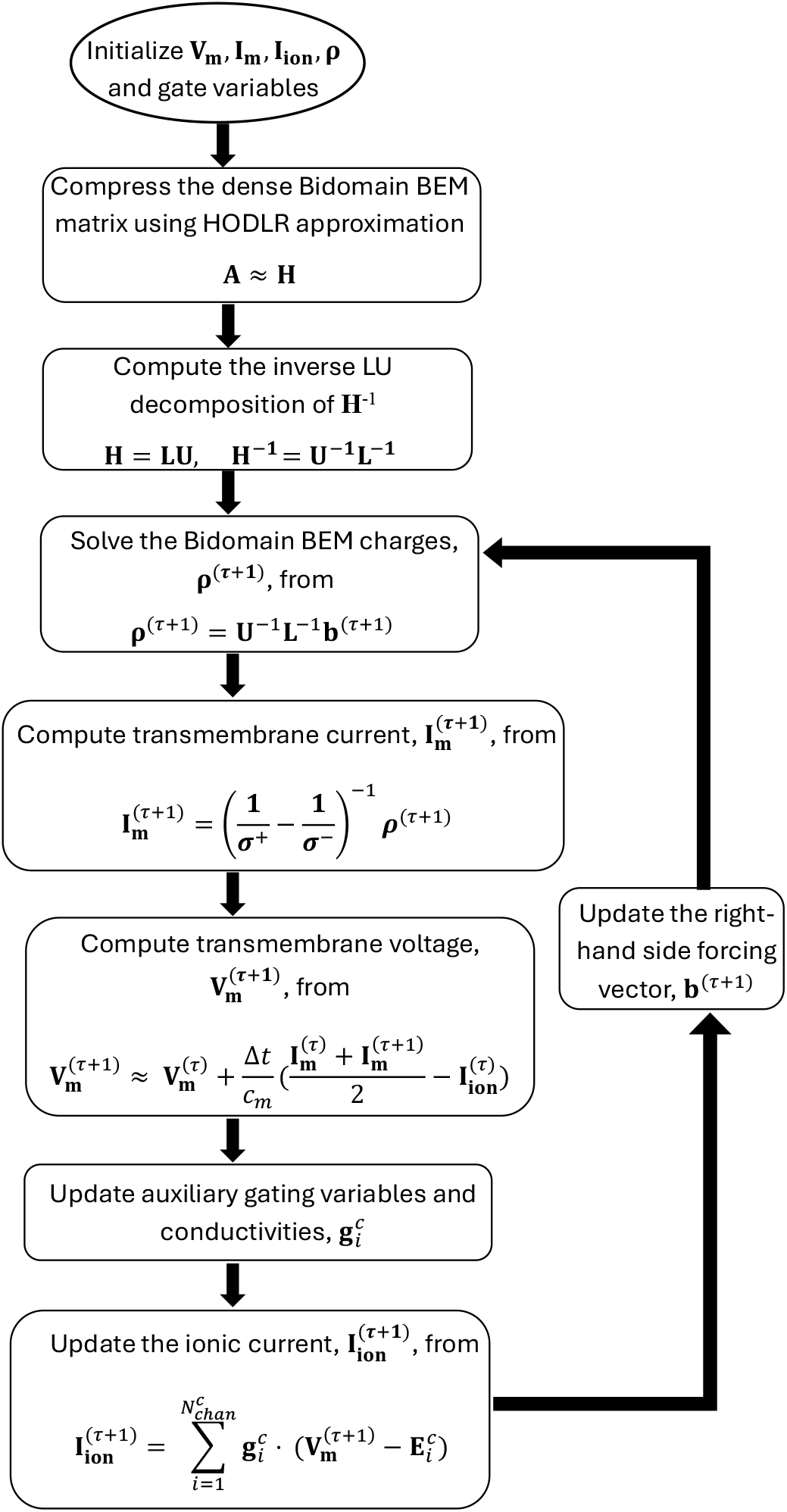
A flowchart of the bidomain BEM algorithm. The algorithm repeats for a user-defined number of time steps.

### E. Meshing a Neuron

A filamentary cable model of the L2/3 pyramidal neuron cell was obtained from the Blue Brain Project [28]. The cable model consists of vertices and the cable path, which is indicated by edges that connect the vertices. The width of each cable segment is indicated for each edge along with its membrane type, which is either soma, axon, apical dendrite, or basal dendrite. To solve the bidomain equation, we must generate a triangle mesh of the neuron surface from the filamentary cable description. This is done by using the algorithm described in [29] with a slight modification when generating the convex hulls. First, a temporary triangle mesh of a circle with a diameter equal to the cable diameter of its corresponding edge is added near each of its ends (Fig. [5B]). Second, individual meshes of edge segments are generated by meshing the convex hull of its two circles. Third, meshes of the vertex junctions are made by meshing the convex hull of all of the nearest circles of coincident edges (Fig. [5C]). Finally, all edge and vertex junction meshes are merged into a single mesh, and repeated triangles (i.e., internal circles) are removed. This results in a watertight mesh (Fig. [5D]). For stability, we modified the algorithm [29] by not generating the convex hull of the vertex junction circle convex hull, but by first mapping all points of circles to a unit sphere centered about its vertex before constructing its convex hull mesh. Fig. [5D] shows an example mesh generated from a 1,387-compartment model of an L2/3 pyramidal cell used in this study. Each triangle is assigned a membrane type.

**Fig. 5:**
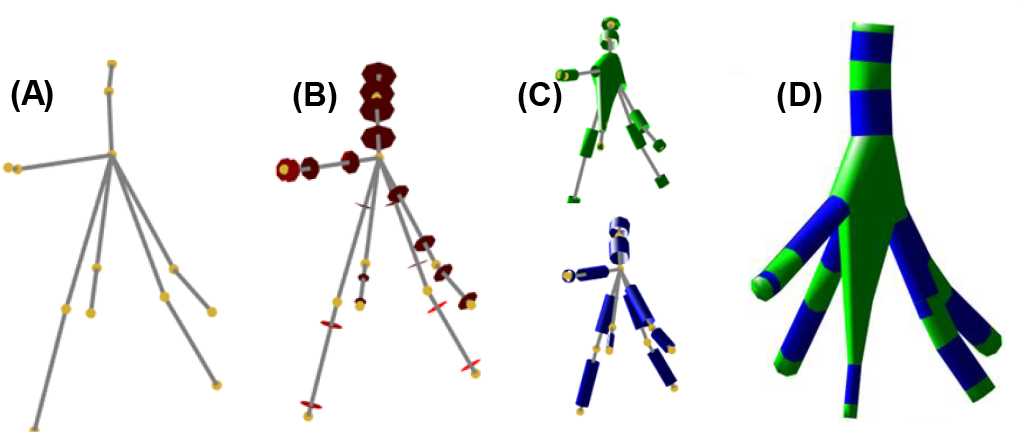
Meshing neuron. (A) node locations and edges of the neuron. (B) Adding a temporary circle triangle mesh (red). (C) Constructing a mesh by connecting all circle meshes of each compartment, and then removing the circle meshes. (D) Merging meshes to construct a surface mesh of the neuron.

## F. Gate Equations

Each compartment has a set of ion channels and equations governing the membrane mechanisms. The equivalent circuit models for each type of compartment are shown in Fig. [6]. Each ionic channel has its conductance and a voltage source that models the electrochemical gradient driving the ion movement between the extracellular and intracellular regions. For example, the axon, soma, apical dendrite and basal dendrite has ion channels with conductivities **g**^*axon*^, **g**^*soma*^, **g**^*apical*^, and **g**^*basal*^, respectively. The corresponding voltage sources are **E**^*axon*^, **E**^*soma*^, **E**^*apical*^, and **E**^*basal*^, respectively. A detailed description of each ion channel is provided in the supplementary file.

**Fig. 6:**
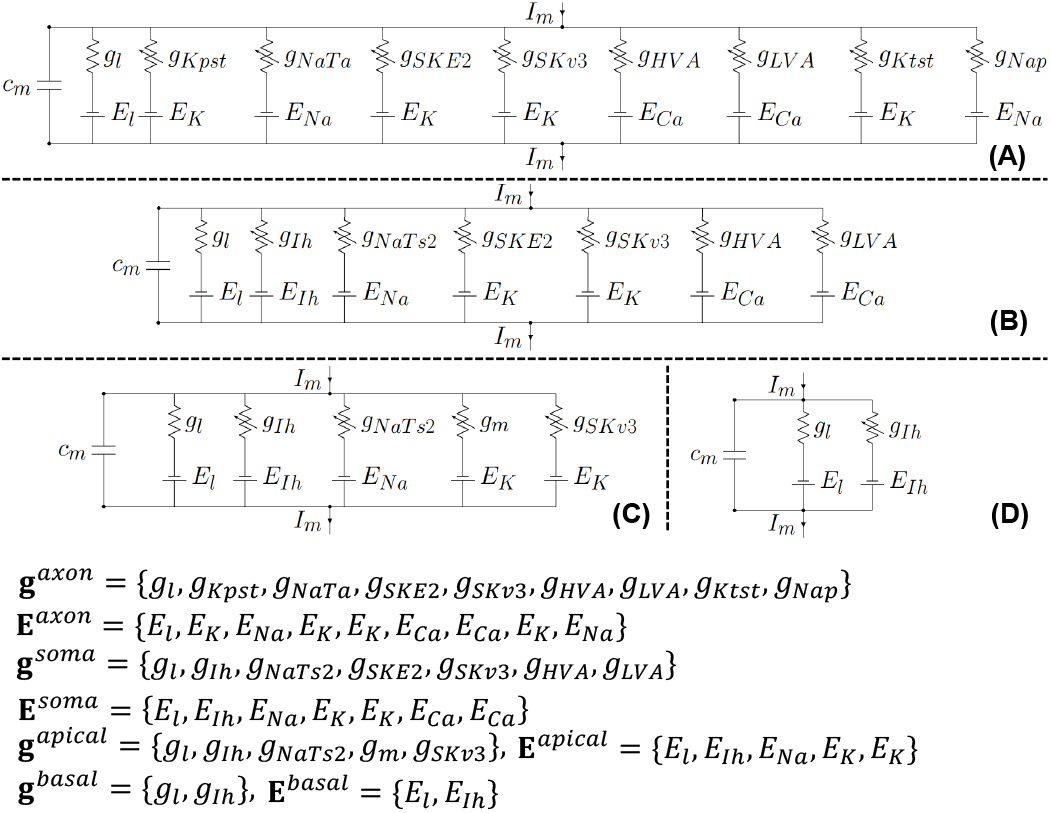
Equivalent circuit of the axon (A), soma (B), apical dendrite (C), and basal dendrite (D). Here, the *g*_{·}_ denote the conductance of the specific ion channel, and the *E*_{·}_ is the voltage source that models the electrochemical gradient driving the specific ion movement through that channel between extracellular and intracellular regions.

The ionic current for each compartment type is

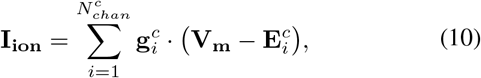

where *c* ∈ {*soma, apical, basal, axon*} and 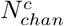 is the number of parallel ion channels in the specific compartment. In the most general sense, the time dependence of each channel is fully described by the following channel parameters 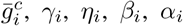, and *τ*_*i*_ summarized in Table IV in the supplementary file. In general, these parameters can be used to compute the elements of conductance vector 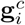 as 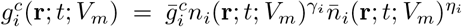, where *n*_*i*_ and 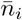 are the probability of the *i*^th^ ion channel being open and close, respectively. Both *n*_*i*_ and 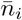 follow the same differential equation

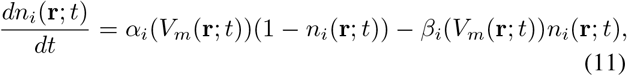

where *α*_*i*_ and *β*_*i*_ are channel dependent parameters. Solving (11), the value of *n*_*i*_ (or,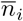) during time-step (*t*+Δ*t*) is found from its value at *t* by the following time-stepping scheme

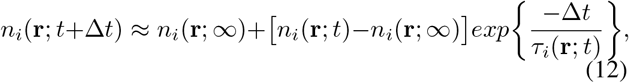

where *n*_*i*_(**r**; ∞) is the limiting probability of the *i*^th^ ion channel being open for a steady value of *V*_*m*_(**r**; *t*) as *t* approaches ∞ and its value for each channel is given in Table I and Table III in the supplementary file. This time-stepping scheme solves the gating probability (11) assuming constant coefficients for the time interval between *t* and *t* + Δ*t*.

### G. Simulation Setup

To model the electrical stimulation of a neuron inside a head compartment, the neuron is submerged inside a cubic bath. The size of the neuron is of the order of 10^−4^m^3^ and the bath is 40 × 40 × 40 mm^3^. The neuron and the bath have conductivities of 1 S/m and 2 S/m. The electrodes are placed on the head surface, two sides of the bath as shown in Fig. [7] with the cathode in green and the anode in red. The electrodes are driven with an external current of amplitude 16 mA, duration of 5 ms (starting at 0.2 ms), and its distributions on both the cathode and anode is assumed uniform. The resulting E-field is nearly uniform and **E**(**r**; *t*) ≈ 80V*/*m along the z-direction (from cathode to anode) for 5 ms starting at 0.2 ms. The bath consists of 2, 168 nodes with 4, 332 triangular facets, and the neuron consists of 16,830 nodes with 33,660 triangular facets; resulting in *N*_*V*_ = 18, 998 nodes and *N*_*T*_ = 37, 992 triangular facets in total. The HODLR matrix consists of *L* = 9 hierarchical levels, where the off-diagonal blocks are computed using ACA with a convergence tolerance of *ϵ*_*H*_ = 10^−7^, respectively. The transmembrane voltage is computed with a temporal resolution of 1 *µ*s and a total of *N*_*t*_ = 40, 000 time steps. The bath and cell are groups using distinct oct-tree hierarchies with 8 levels each to ensure that the first level separates the cell and bath into one group. This partition enables us to determine the strength and rank of the coupling between the cell and the bath. The equilibrium potential for the neuron membrane is set to -74.5 mV.

**Fig. 7:**
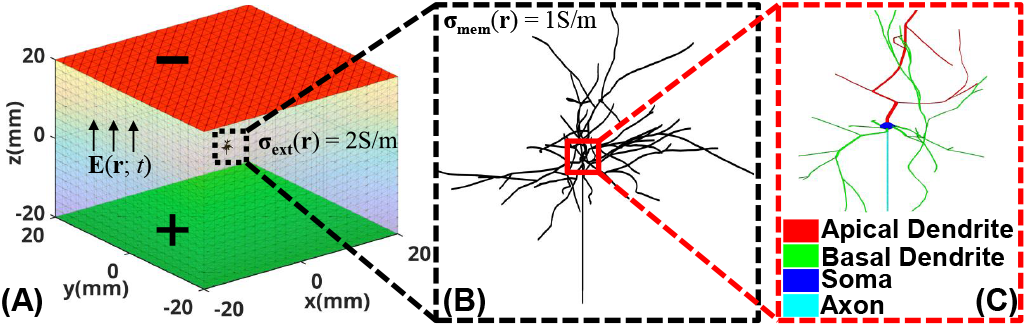
The L2/3 pyramidal neuron is placed within an inhomogeneous medium. The intracellular and the extracellular spaces have conductivities of 1 S/m and 2 S/m, respectively. The cell has four compartment types (soma, axon, apical dendrite, and basal dendrite).

## III. Results

## A. HODLR Matrix Ranks

A compressed form of the HODLR matrix is shown in Fig. [8], using 8 levels for both membrane nodes and bath nodes with *L* = 9. The charges on the neuron membrane generate an E-field that can be decomposed into a relatively small number of basis vectors—i.e., the interaction between membrane charges and evaluation points on the bath surface is of low rank. For example, the neuron-bath and bath-neuron couplings are represented using ranks of only 3 and 9, respectively.

**Fig. 8:**
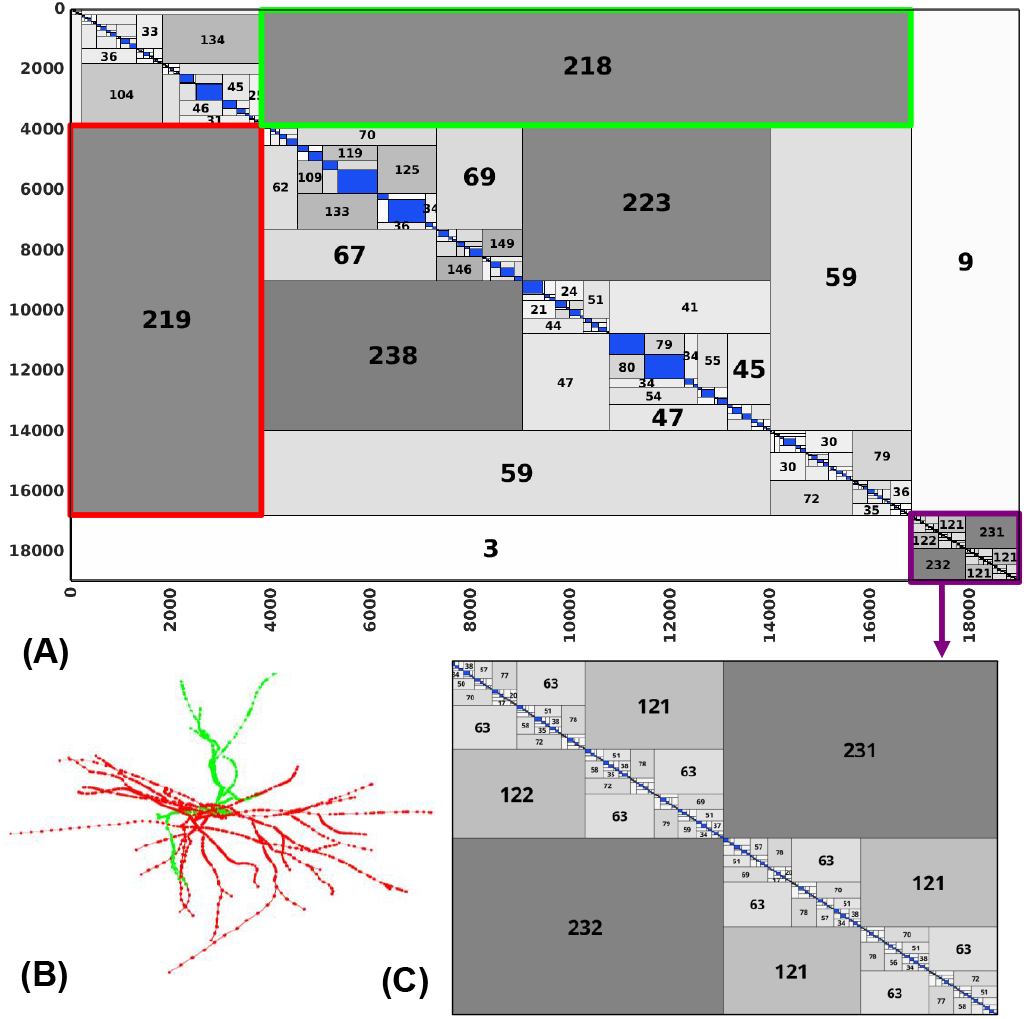
(A) HODLR representation of the BEM matrix (dense blocks in blue). (B) Two sets of membrane nodes with weak interaction, shown in green and red blocks (3, 845 × 12, 986) in (A), require only 219 and 218 ranks. (C) HODLR subblock for the interaction of charges on the bath boundary.

Additionally, E-fields generated by charges at different group locations are also of low rank. For instance, the green and red submatrices in Fig. [8] correspond to interactions between the bottom and top halves of the cell membrane (Fig. [8B]). These submatrices have sizes of 3, 845 × 12, 986 and ranks of 219 and 218, resulting in a compression ratio of 2980. Fig. [8A] highlights the inadmissible dense blocks in blue. The submatrix of the bath is less compressible than that of the membrane (Fig. [8C]), likely because the membrane is a wire-like structure with fewer unknowns per unit volume compared to the solid bath boundary. The maximum rank required across all sub-blocks is 238.

### B. Compression Ratio and Computational Run-time

All benchmarking results were obtained using an AMD EPYC 7502 32-core processor (2495.535 MHz) CPU. Fig. [9A] shows the required computational memory as a function of the number of neurons under stimulation, along with the corresponding compression ratio, defined as the ratio of memory required by the uncompressed BEM matrix to that of the HODLR-compressed version. Fig. [9B] displays the setup time for HODLR compression.

**Fig. 9:**
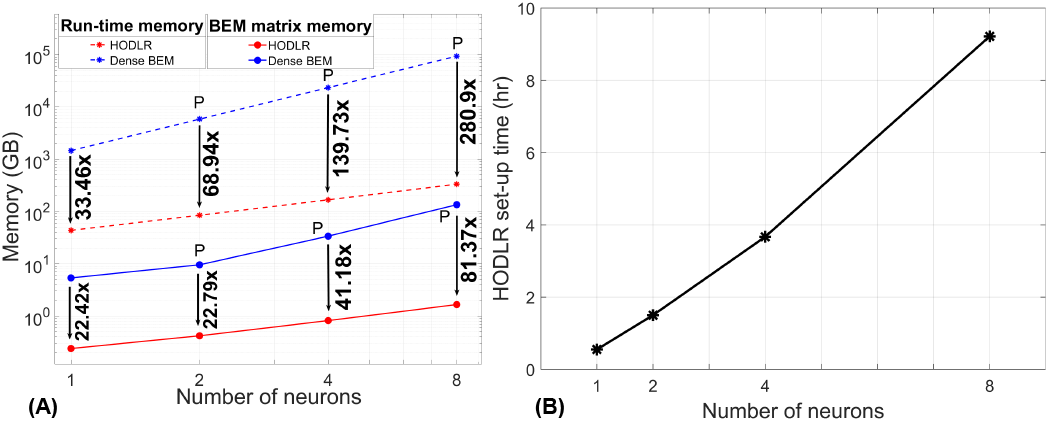
(A) Computational run-time memory and the BEM matrix memory for the HODLR compression and the uncompressed dense BEM matrix as a function of the number of neurons. ‘P’ denotes the entries computed with the interpolation technique. Memory compression is shown as bold text. (B) Set-up time required by the HODLR compression method with respect to the number of neurons.

For a single neuron (18,998 nodes and 37,992 triangular facets), computing the full dense BEM matrix requires 5.38 GB of memory, while the compressed HODLR matrix requires only 0.24 GB, yielding a 22.4× compression. The setup times for the uncompressed and HODLR matrices are 77.92 and 32.88 minutes, respectively. Over 40,000 time steps, the transmembrane voltage computation takes 191.08 minutes (0.29 seconds/step) using the dense BEM matrix and only 19.6 minutes (0.03 seconds/step) using the HODLR matrix, resulting in a 10× speedup.

For a group of 8 neurons (136,808 nodes and 273,612 triangular facets), the HODLR matrix requires just 1.66 GB of memory compared to 135.07 GB for the dense BEM matrix—an 81.37× reduction. During the setup phase, peak RAM usage is 332.78 GB for HODLR and 93,478 GB for the dense BEM matrix, corresponding to a 280.90× compression. The total setup time for the 8-neuron configuration is 9.22 hours (Fig. [9B]).

## C. Accuracy Benchmarks

Fig. [10] compares the predicted membrane voltage at representative points along the axon, soma, apical dendrite, and basal dendrite, as computed by the cable equation-based solver, the uncompressed BEM, and the HODLR-compressed

**Fig. 10:**
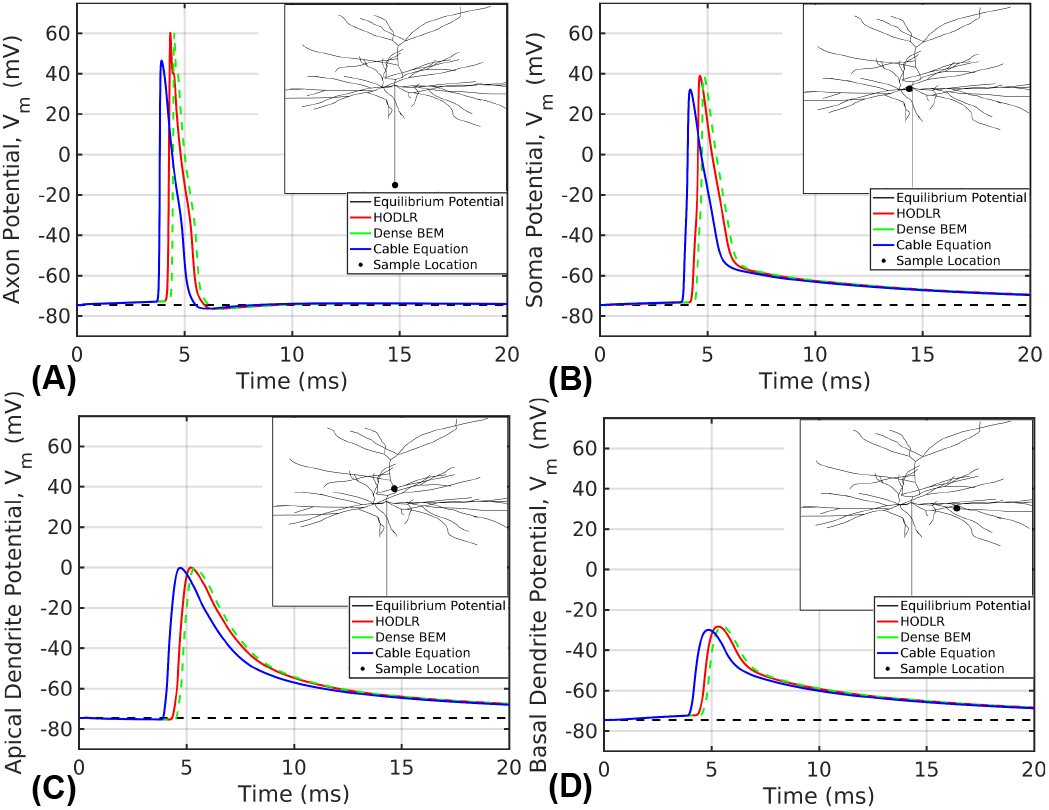
Propagation of action potential (V_m_) through the axon (A), soma (B), apical dendrite (C) and basal dendrite (D) computed from the uncompressed dense BEM matrix (green), the HODLR compression (red) and the cable equation approach (blue). The inset shows the corresponding spatial sample locations (black dot).

The action potentials are generated with an The uncompressed and HODLR-compressed BEM results are virtually indistinguishable, confirming the fidelity of the HODLR compression. However, the cable equation-based solution shows an earlier activation, approximately 0.40 ms ahead of the BEM results. At the axon (and soma), the cable equation also predicts a lower peak membrane voltage of 60.16 mV (and 38.98 mV), compared to 46.44 mV (and 32.14 mV) for both BEM-based solvers. This discrepancy likely stems from the simplified activation function used in cable models, as opposed to the more detailed electric field coupling captured by BEM. The action potential initiates at the distal axon and propagates through the axon, soma, and dendritic structures. The propagation speed is primarily influenced by the local cross-sectional area of the neuron. The consistent 0.40 ms delay across all sample locations in Fig. [10] suggests that the cable and BEM models exhibit the same propagation speed, despite the timing offset.

Fig. [11] provides additional detail at the axonal sample site. It shows the membrane voltage (V_m_), transmembrane current (I_m_), total channel conductance (Σ_*i*_ g_*i*_), membrane charge density (*ρ*), and total ionic currents (I_ion_). Notably, the transmembrane current, total ionic current, and membrane charge density each undergo two polarity reversals—once at action potential initiation and again at the peak membrane voltage. Individual ionic channel contributions are shown in Fig. [11A]. Some channels, such as NaTa, SKv3, and Kpst, contribute significantly to the total current (up to 708 A), while others, like Ktst and Nap, exhibit much smaller currents (up to 16 A). Calcium channels, the SKE2 channel, and leakage currents contribute negligibly, up to 1 A.

**Fig. 11:**
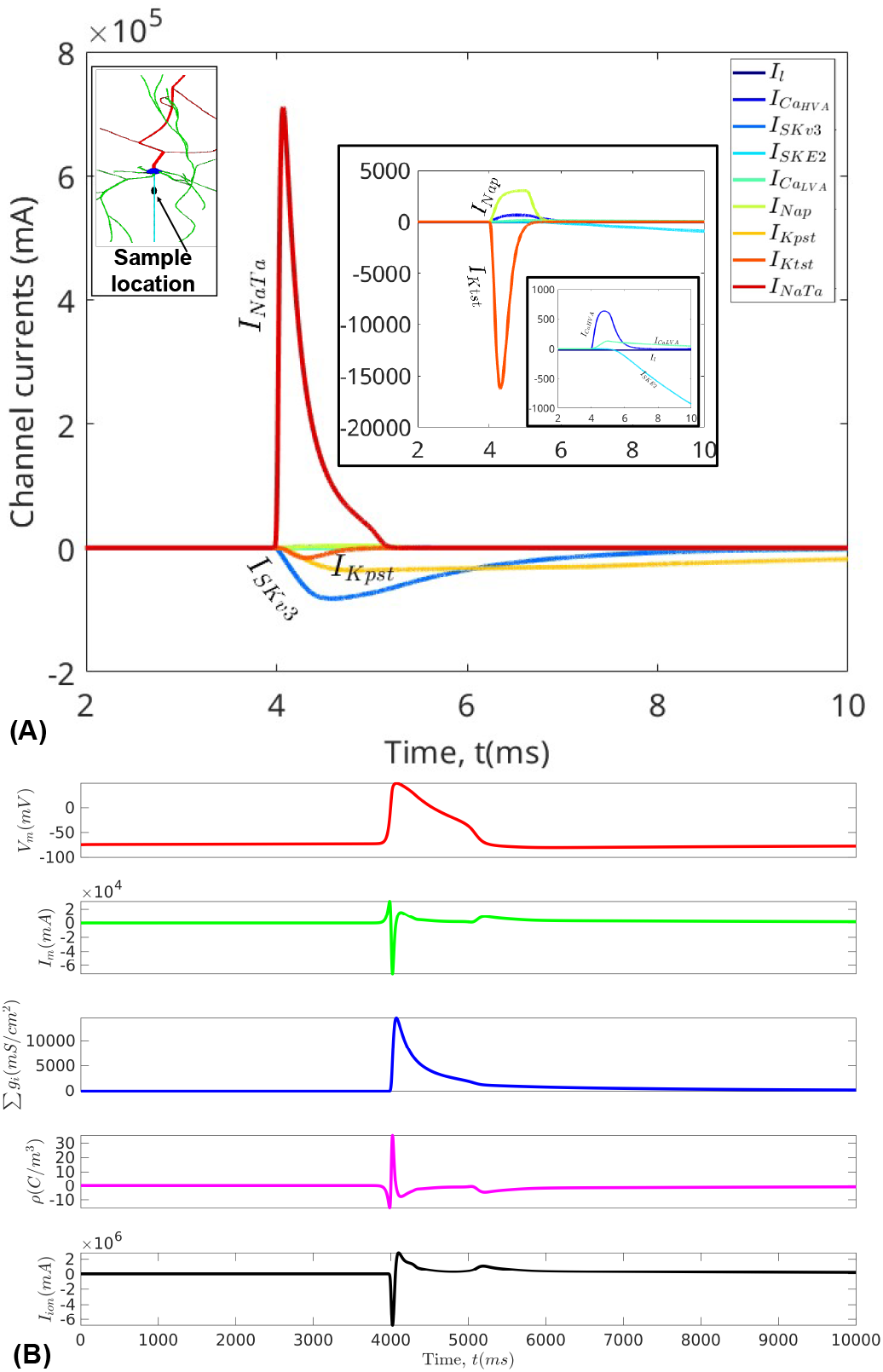
(A) For the sample coordinate along the axon (inset), individual ionic channel currents are shown; (B) transmembrane voltage (V_m_), transmembrane current (I_m_), total channel conductance (Σ_*i*_ g_*i*_), membrane charge density (*ρ*) and total ionic currents (I_ion_) for the sample location in (A).

Fig. [12] illustrates the spatiotemporal propagation of the action potential across the neuronal membrane at successive time steps. The action potential initiates at the distal tip of the axon (Fig. [12A]), where the external electric field generated by the electrodes is optimally aligned with the axonal orientation. The depolarization then propagates upward through the soma (Fig. [12](B–C)]), before attenuating as it spreads into the dendritic branches (Fig. [12](D–F)]). This behavior closely aligns with predictions made by cable equation-based models and reflects known biophysical dynamics of action potential propagation in neurons.

**Fig. 12:**
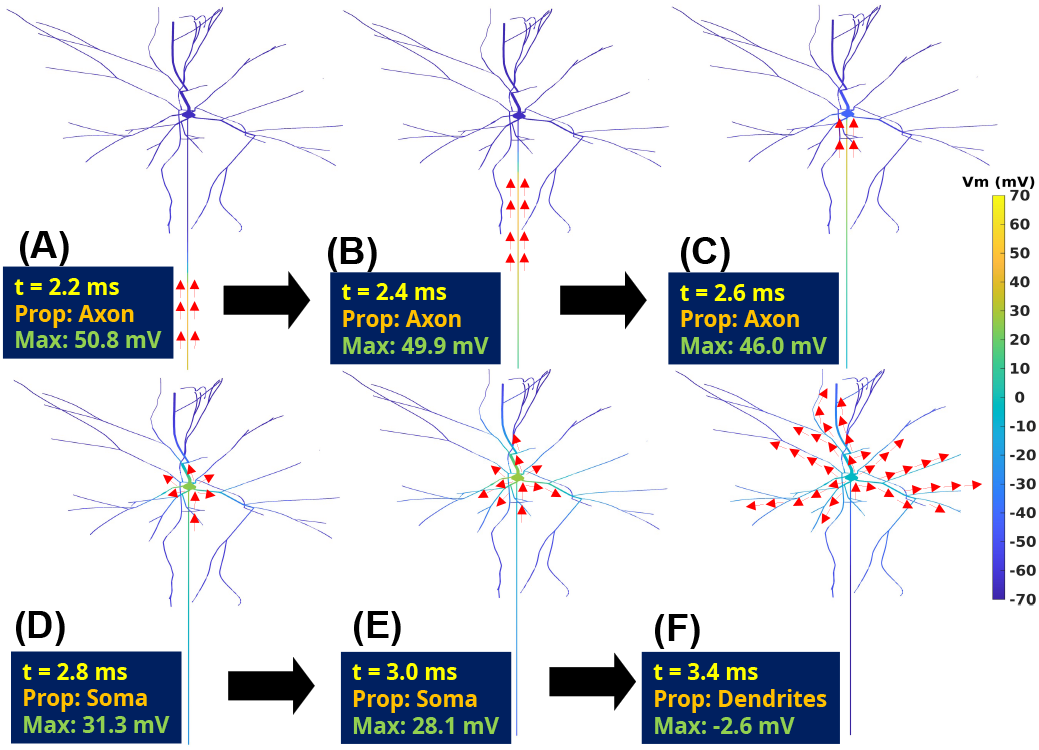
The action potential starts propagating from one end of the axon (A) and traverses through the axon towards the soma (B-C). Next, it propagates along the dendrites to nearby neurons (D-F). The insets show the time stepping (*t*), propagating compartment (*prop*), and the maximum amplitude of the action potential. The red arrow follows the maximum transmembrane voltage.

### D. Activation threshold of a realistic neuron

In this section, we compare the activation thresholds predicted by the cable equation and the BEM approach for a realistic neuron model (Fig. [13]) for an equilibrium potential of -74.5 mV.

**Fig. 13:**
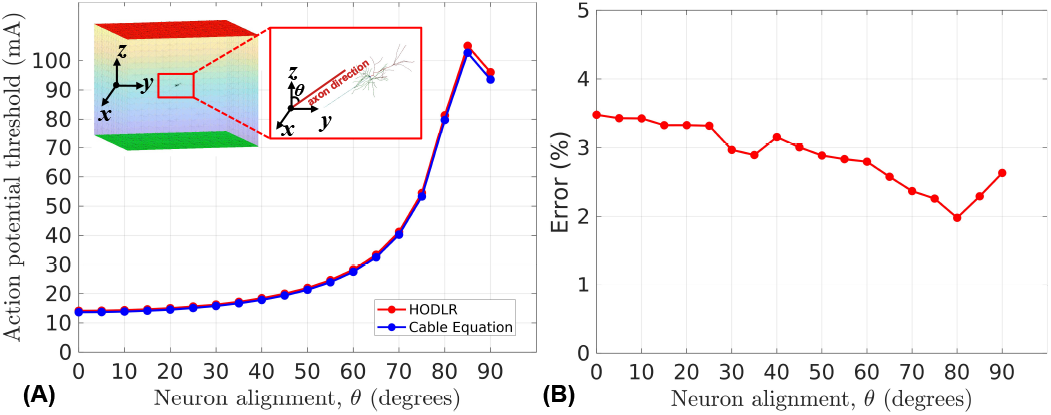
Effect of neuron spatial orientation on the action potential (A). The action potential threshold increases as a function of the angle (*θ*) between the incident field and the axon for both the BEM and the cable-equation approach. (B) The relative error of the BEM threshold with respect to the cable-equation threshold increases as a function of the neuron spatial orientation (*θ*).

The orientation angle *θ* between the *z*-axis (i.e., the direction of the external electric field) and the neuron’s axon is varied as a parameter (Fig. [13A]). When the neuron is fully aligned with the E-field (*θ* = 0°), the activation thresholds are 14.16 mA for the BEM approach and 13.68 mA for the cable equation. When the neuron is oriented transversely to the E-field (*θ* = 90°), the thresholds are 96.03 mA for BEM and 93.57 mA for the cable model. This corresponds to BEM-based thresholds that are 3.48% and 2.63% higher than those predicted by the cable equation for *θ* = 0° and *θ* = 90°, respectively (Fig. [13B]). Across the range of orientations, the BEM approach consistently yields slightly higher activation thresholds. The smallest difference is 1.98% at *θ* = 80°, and the largest is 3.48% at *θ* = 0°.

### E. Multiple Neurons

In this section, we analyze both compression efficiency and action potential behavior in simulations involving multiple neurons. Fig. [14A] illustrates a scenario in which a second neuron (N_2_) gradually moves closer to a target neuron (N_1_). The corresponding HODLR-compressed matrix is shown in Fig. [14B]. The E-field that the neurons generate on each other is highly compressible, requiring a rank of only 17 or 18. Fig. [14C] shows how the presence of N_2_ affects the activation threshold of N_1_ as N_2_ approaches either along the x-direction or the z-direction. The results indicate that the action potential threshold of N_1_ is only minimally affected by N_2_’s proximity. However, the threshold is slightly lower when N_2_ approaches along the z-direction compared to the x-direction. This may be because the external coil E-field impinges directly on N_2_ when it moves along x, while N_1_ partially shields it when it moves along z.

**Fig. 14:**
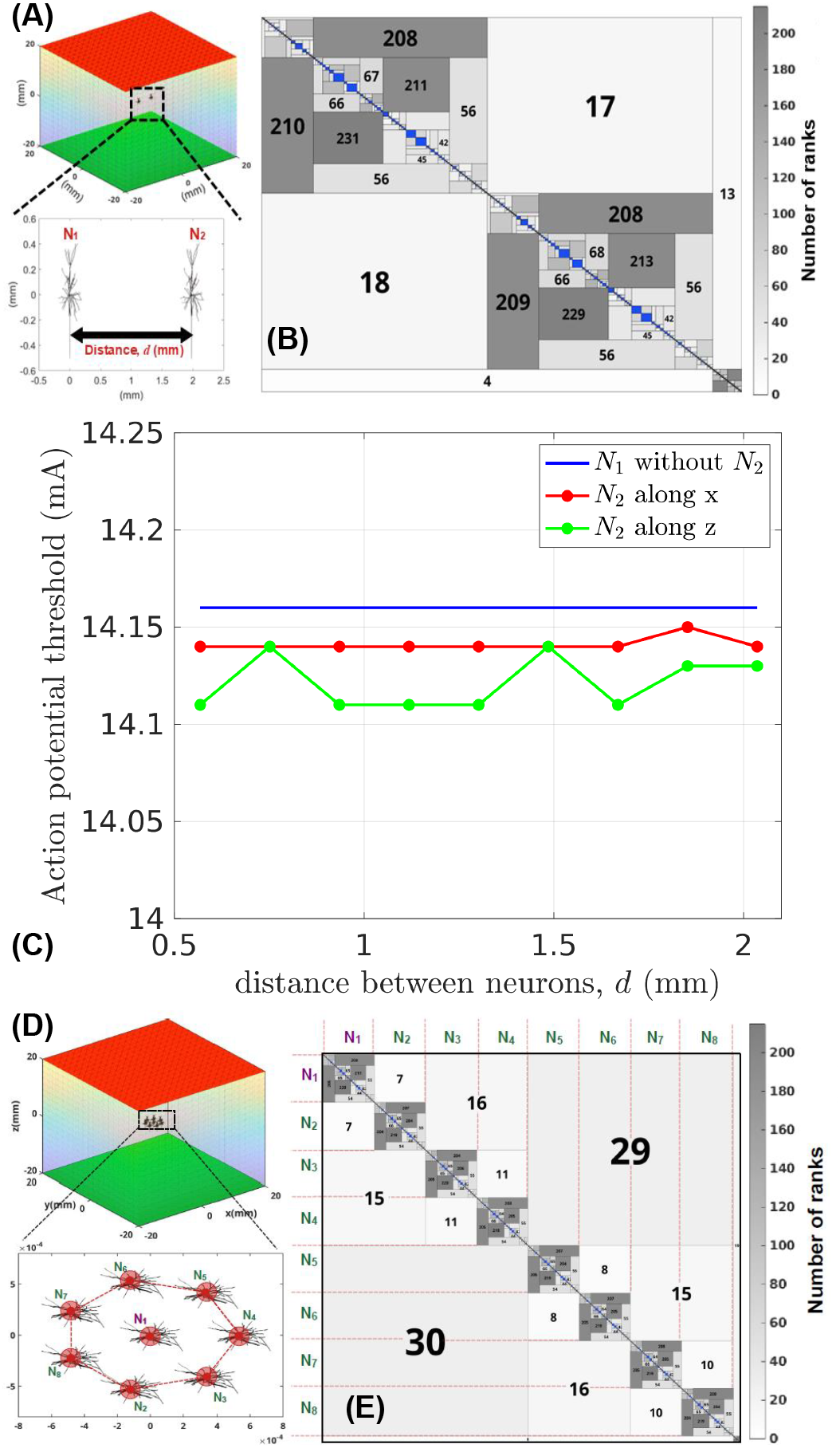
(A) Two neurons at a distance, d, under the stimulation inside the bath. (B) HODLR compressed matrix for the two-neuron scenario. (C) The action potential threshold of the observed neuron (N_1_) is lower when the second neuron (N_2_) moves along the z-direction (parallel to the stimulating E-field) than along the x-direction. (D) A group of 7 neurons (N_2_-N_8_) surrounds an observed neuron (N_1_) in a regular heptagon pattern. (E) The resultant HODLR matrix of the group of 8 neurons.

A similar analysis is performed for a group of 8 neurons (Fig. [14](D–E)]). Here, the observed neuron N_1_ is surrounded by 7 other neurons (N_2_–N_8_), placed at the vertices of a regular heptagon (Fig. [14D]), with all neurons aligned along the direction of the stimulating E-field. The resulting HODLR matrix (Fig. [14E]) again shows highly compressible interactions between the neurons. As in the two-neuron case, the action potentials of all neurons remain unaffected by the presence of nearby cells.

These results demonstrate that our HODLR-based BEM framework effectively handles multi-neuron simulations, maintaining accuracy while offering significant compression.

## IV. Discussion

Scalable bidomain modeling of neuron stimulation by a device-induced E-field using the BEM approach has been introduced by leveraging fast direct solvers. The computation and storage of the dense BEM matrix for a realistic neuron are computationally intractable. Additionally, updating the neuron membrane voltage at every time step requires recomputing the dense BEM matrix, limiting the applicability of the standard bidomain method. By utilizing the hierarchical off-diagonal low-rank (HODLR) form, the dense BEM matrix was precomputed and stored. The HODLR form reveals that interactions between charges on the membrane and the head surface, as well as interactions between charges on the same neuron, are highly compressible. An 8-level HODLR structure is sufficient to compute the BEM matrix without compromising the accuracy of action potential generation. A multi-neuron simulation scenario shows that the action potential of any given neuron is negligibly affected by nearby neurons.

Validation results indicate that using an 8-level hierarchical binary partitioning of the membrane nodes compresses the dense BEM matrix by 22.42 times for an L2/3 pyramidal neuron. Additionally, runtime memory is reduced by 33.46 times for the same neuron. With our existing hardware resources, simulations of up to 8 neurons show a BEM matrix compression of 81.37 times and a runtime memory reduction of 280.9 times. Moreover, it takes only 0.55 hours and 9.22 hours to precompute and store the BEM matrix in HODLR form for a single neuron and a group of 8 neurons, respectively. Our results show that by compressing the inverse of the BEM system of equations, it is feasible to solve for multiple cells using the bidomain model. The runtime memory of the solvers can likely be further reduced by performing computations in batches or, for direct solvers, by avoiding recursive implementations. For HODLR, there is a three-order difference in required storage between the runtime processes and the actual BEM matrix. This suggests that, by reducing runtime memory usage, thousands of cells could be modeled on a standard computing cluster. Computational time increases linearly with the number of cells, but this can be improved by parallelizing the generation of the HODLR matrices. Finally, the coupling ranks between cells and the surrounding medium, or between neighboring cells, tend to be low. This indicates potential for further improvement through matrix factorization using Schur complements, which could eliminate the need to solve for the surrounding media boundary at every time step. With these improvements, using HODLR solvers to analyze networks of thousands of neurons using BEM could become practical.

All results presented here indicate that cable solvers can capture the key features of stimulation by a device-induced E-field. The BEM introduces subtle differences, likely due to the inclusion of geometric factors. Incorporating elliptical crosssections for cell segments, more realistic representations of the soma, and the inclusion of dendritic spines could lead to further deviations from the cable model, and these effects warrant further study. As shown in our previous work [10], both the cable equation and BEM demonstrate a similar relationship between activation threshold and cell angle, suggesting that the key features of activation are preserved in the simplified cable model, with differences arising from more nuanced effects.

Multi-neuron scenarios have recently been shown to yield different activation thresholds than single-cell simulations [10], [30]. These scenarios cannot be accurately studied using standard cable approaches, which assume that the cell’s activity and boundaries do not distort the device-induced E-fields. This work represents a significant step toward enabling the practical simulation of multi-neuron scenarios while preserving full geometric representations of cells and more accurately predicting the effects of E-fields on neural networks.

## V. Conclusion

By leveraging direct solvers, we presented a systematic approach to compress the dense bidomain BEM matrix using the HODLR form, enabling full coupling of the neuron, the inhomogeneous medium, and the stimulating device for a realistic L2/3 rat pyramidal neuron. Compression and storage of the dense BEM matrix take only 0.55 hours, achieving compression ratios of 22.42 times for the matrix itself and 33.46 times for maximum runtime memory. Future work will explore the influence of geometric factors on stimulation and expand to multi-cell scenarios.

