## Supplementary_File for "Modeling Pyramidal Neurons Using Bidomain BEM and Hierarchical Matrix Approximation"

### VI. Supplementary File

#### S1. A 3-level HODLR Form

Fig. [15] shows a 3-level HODLR matrix, and Fig. [16] shows its LU decomposition at each level.

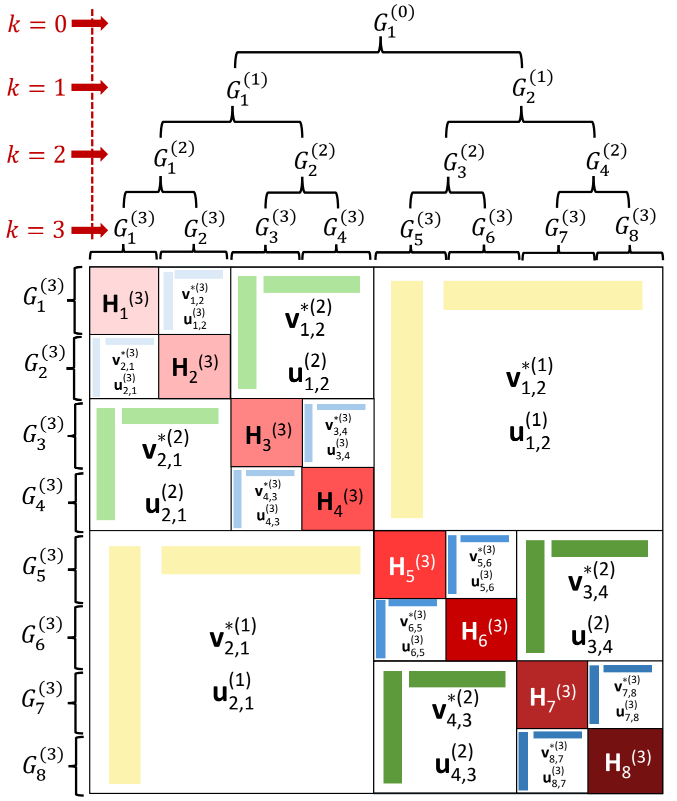


Fig. 15: HODLR matrix for 3 levels of hierarchical partitioning. The $k^{\text{th}}$ level has $2^k$ groups of nodes. The off-diagonal admissible blocks are computed using a low-rank approximation technique. At the last level ($k = 3$), the diagonal inadmissible dense blocks $[\mathbf{H_1^{(3)}}, \mathbf{H_2^{(3)}}, \ldots, \mathbf{H_8^{(3)}}]$ are shown in solid red colors.

#### S2. Ion Channels

Table I, II, and III summarize different ion channels, membrane capacitance, and their steady-state conductance with reverse potential, respectively.

#### S3. Determining Ca ion concentration ($\mathrm{Ca}^c(\mathbf{r};t)$) and Ca equilibrium potential ($\mathrm{E_{Ca}}^c(\mathbf{r};t)$)

$$
\begin{aligned}
\mathrm{E_{Ca}}^c(\mathbf{r};t+\Delta t) &= \frac{\mathrm{RT}}{2\mathrm{F}} log\left(\frac{2}{\mathrm{Ca}^c(\mathbf{r};t+\Delta t)}\right), \\
\mathrm{Ca}^c(\mathbf{r};t+\Delta t) &= \mathrm{Ca}^c(\mathbf{r};\infty) \\
&\quad + \left[\mathrm{Ca}^c(\mathbf{r};t) - \mathrm{Ca}^c(\mathbf{r};\infty)\right]\cdot \\
&\qquad exp\{-\frac{\Delta t}{\tau^c_{\mathrm{Ca}}(\mathbf{r};t)}\}, \\
\mathrm{Ca}^c(\mathbf{r};\infty) &= -10000\mathrm{d}\frac{\mathrm{I_{Ca}}^c(\mathbf{r};t)\Gamma}{2\mathrm{FD}} + 10^{-4}, \\
\mathrm{I_{Ca}}^c(\mathbf{r};t) &= \sum_{i\in\{\mathrm{Ca_{HVA}},\mathrm{Ca_{LVA}}\}} \bar{\mathrm{g}}^c_i \cdot [\mathrm{n}_i(\mathbf{r};t)]^2 \cdot \bar{\mathrm{n}}_i(\mathbf{r};t)\cdot \\
&\qquad \left[\mathrm{V_m}(\mathbf{r};t) - \mathrm{E_{Ca}}^c(\mathbf{r};t)\right],
\end{aligned} \tag{13}
$$

where $\mathrm{d} = 287.198731\ \mathrm{ms}^{-1}$ is the decay rate of removal of Ca ion, $\mathrm{F} = 96485.33212\ \mathrm{Cmol}^{-1}$ is the Faraday constant, $\mathrm{D} = 0.1\ \mu m$ is the depth of shell, $\tau_{\mathrm{Ca}} = 287.198731\ \mathrm{ms}^{-1}$ is the time constant, $\Gamma = 0.002910$ is the percent of free Ca ion (not buffered), $\mathrm{R} = 8.3144598\ \mathrm{Jmol}^{-1}\mathrm{K}^{-1}$ is the gas constant, $\mathrm{T} = 34^o\mathrm{C}$ (307.15 K) is the absolute temperature. The values of $\bar{\mathrm{g}}^c_i$, $\mathrm{M}_i(\mathbf{r};t)$ for a specific compartment $c$ are found from Table III and Table IV, respectively, and by using Eq. (12).

#### S4. Gating Variables

Table IV summarizes the formula for computing the probability of an ion channel being open or closed.

#### S5. Effect of External Bath and Hierarchical Tree Structure

In this section, we analyze the effect of the dimension of the bath ($n \times n \times n$) and the number of HODLR hierarchical levels ($l$) on the computational memory, maximum HODLR rank, delay in the initialization of action potential, and the preprocessing time for a single neuron under stimulation (Fig. [17]). The required memory, the maximum rank in the HODLR matrix, and the preprocessing time increase as a function of the bath dimension shown in Fig. [17A], Fig. [17B], and Fig. [17D], respectively. The bath dimension negligibly affects the initialization of the action potential (Fig. [17C]). On the other hand, the computational memory and the maximum HODLR rank decrease as a function of HODLR hierarchical levels, shown in Fig. [17A] and Fig. [17B], respectively. The HODLR hierarchical levels negligibly affect the initialization of action potential (Fig. [17C]) and the preprocessing time (Fig. [17D]). Neither the bath dimension nor the tree levels affect the initialization and the total computational time of transmembrane voltage.

#### S6. Effect of neuron spatial angle of rotation on action potential initiation

Fig. [18] shows the variation of action potential initiation time as a function of the spatial alignment of the neuron for different stimulating currents. As the angle between the

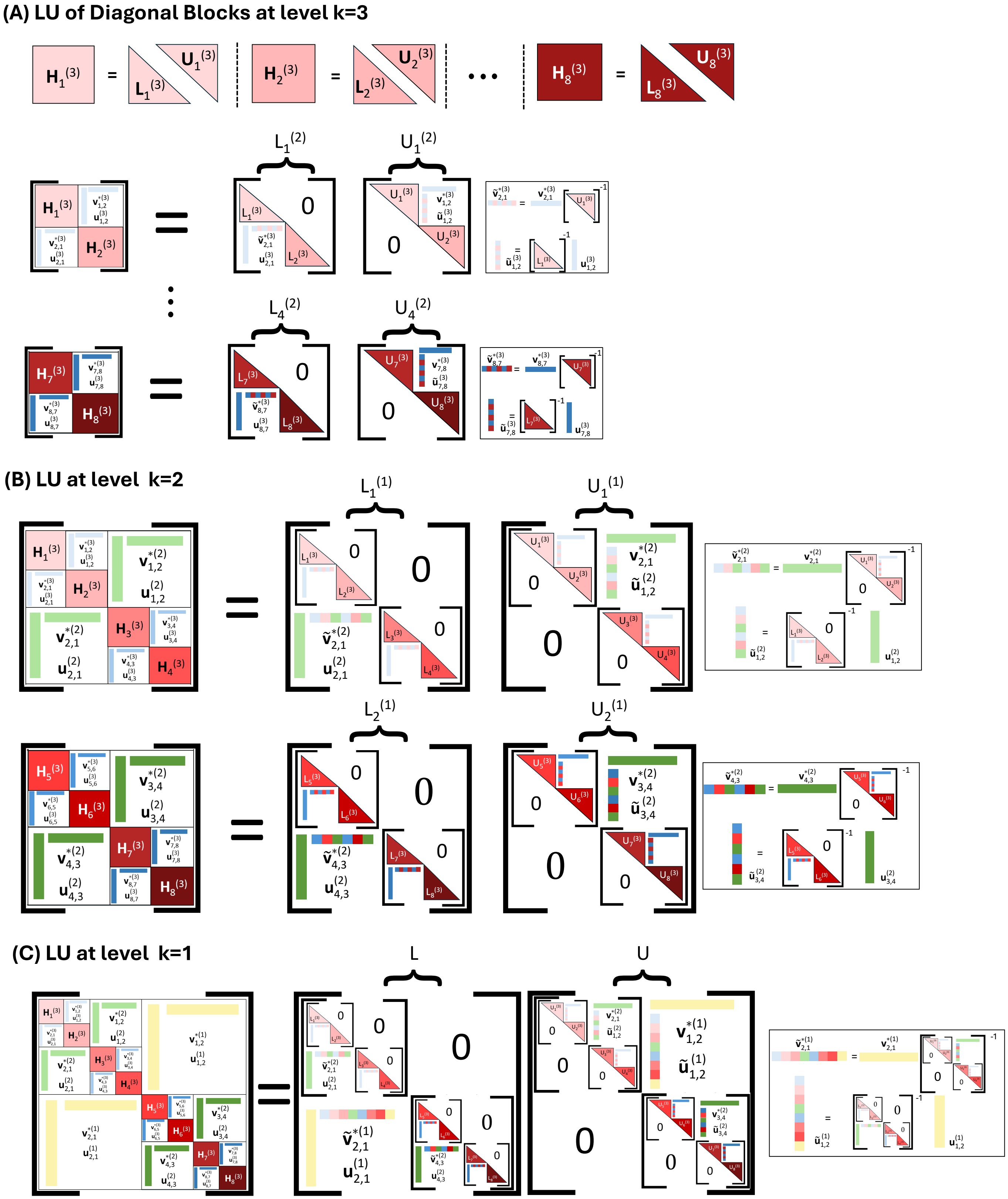


Fig. 16: LU decomposition of the HODLR matrix. (A) Diagonal inadmissible blocks at level $k = 3$ are factorized using direct LU decomposition. (B-C) LU decomposition at level $k = 2$ and $k = 1$, respectively.

TABLE I: Ion channel information

| Channel name | Description | Source |
| --- | --- | --- |
| $SK_{v3}$ | Shaw-related potassium channel family in rat brain. | Jens et al. [31] |
| $SK_{E2}$ | Small-conductance, Calcium-activated Potassium channels from mammalian brain. | Köhler et al. [32] |
| Nap | sustained, or "persistent," $Na^+$ current channel producing subthreshold oscillatory activity in entorhinal cortex layer-II principal neurons. | Jacopo et al. [33] |
| $K_{pst}$ | The persistent component of the $K^+$ current. | Korngreen et al. [34] |
| $K_{tst}$ | The transient component of the $K^+$ current. | Korngreen et al. [34] |
| NaTa | Modified $Na^+$ channel conductance due to the axonal biophysical properties in the axon hillock, or initial segment. | Colbert et al. [35] |
| Ih | hyperpolarization-activated cation current channel. | Kole et al. [36] |
| Im | M-currents in bullfrog sympathetic neurons. | Adams et al. [37] |
| NaTs2 | Somatic $Na^+$ channel but shifted both activation/inactivation of the channel by 6 mV from the NaTa channel. | Colbert et al. [35] |
| $Ca_{HVA}$ | High-voltage-activated calcium channel. | Reuveni et al. [38] |
| $Ca_{LVA}$ | Low-voltage-activated calcium channel in hippocampal CA3 pyramidal neurons. | Avery et al. [39] and $\tau$ from Randall et al. [40] |

TABLE II: Membrane capacitance

| Membrane type | Capacitance ($\mu F cm^{-2}$) |
| --- | --- |
| Axon | 1 |
| Soma | 1 |
| Apical Dendrite | 2 |
| Basal Dendrite | 2 |

TABLE III: Ion channel conductance and equivalent potentials

| Ion channel | channel conductance notation | Maximum channel conductance, $\bar{g}_i^{\{c\}}$ (mS/cm$^2$) | | | | Equivalent potential, $E_i^{\{c\}}$ (mV) |
| --- | --- | --- | --- | --- | --- | --- |
| | | Axon | Soma | Apical Dendrite | Basal Dendrite | |
| $SK_{v3}$ | $g_{SKv3}(V_m;t)$ | 10219.45 | 3034.72 | 42.26 | - | $E_K = -85.0$ |
| $SK_{E2}$ | $g_{SKE2}(V_m;t)$ | 71.04 | 84.07 | - | - | $E_K = -85.0$ |
| Nap | $g_{Nap}(V_m;t)$ | 68.27 | - | - | - | $E_{Na} = 50.0$ |
| $K_{pst}$ | $g_{Kpst}(V_m;t)$ | 9735.38 | - | - | - | $E_K = -85.0$ |
| $K_{tst}$ | $g_{Ktst}(V_m;t)$ | 892.59 | - | - | - | $E_K = -85.0$ |
| NaTa | $g_{NaTa}(V_m;t)$ | 31379.68 | - | - | - | $E_{Na} = -50.0$ |
| Ih | $g_{Ih}(V_m;t)$ | - | 0.8 | 0.8 | 0.8 | $E_{Ih} = -45.0$ |
| Im | $g_m(V_m;t)$ | - | - | 1.43 | - | $E_m = -85.0$ |
| NaTs2 | $g_{NaTs2}(V_m;t)$ | - | 9839.55 | 261.45 | - | $E_{Na} = -50.0$ |
| $Ca_{HVA}$ | $g_{HVA}(V_m;t)$ | 9.9 | 9.9 | - | - | ${E_{Ca}}^{\{c\}}(\mathbf{r};t)$ |
| $Ca_{LVA}$ | $g_{LVA}(V_m;t)$ | 87.52 | 3.33 | - | - | ${E_{Ca}}^{\{c\}}(\mathbf{r};t)$ |
| Leakage | $g_l$ | 0.3 | 0.3 | 0.3 | 0.3 | $E_l = -75.0$ |

incident field and the neuron increases, it requires higher stimulating currents to initiate an action potential. For the same alignment, the lower stimulating current results in a slower initiation than that of the higher stimulation current.

#### S7. Comparison with the fast multipole method

An alternate choice of computing the surface charge densities ($\rho(\mathbf{r};t)$) is through the fast multipole method (FMM). But FMM does not invert the dense BEM matrix ($\mathbf{A}$). It requires using an iterative solver to solve for the charges. As shown in Fig. 19, for the first few time steps (until 38$^{th}$ iteration), the FMM computational time is lower than that of HODLR compression. However, across 40,000 time steps, the FMM requires 511 hours (due to the iterative solver at every time step), whereas the HODLR compression requires only 0.88 hours (simple matrix-vector multiplication at every time step).

TABLE IV: Gating variables for different active ion channels in the neuron membrane

| Ion channel | Gating variables | Channel open probability, $\left[\mathrm{n}(\mathbf{r};t)\right]^{\gamma} \cdot \left[\overline{\mathrm{n}}(\mathbf{r};t)\right]^{\eta}$ |
|---|---|---|
| $\mathrm{Ca_{HVA}}$ | $\alpha_{\mathrm{n}}(\mathbf{r};t) = \frac{0.055(\mathrm{V_m}(\mathbf{r};t)+27)}{1-e^{-\frac{\mathrm{V_m}(\mathbf{r};t)+27}{3.8}}}$, $\beta_{\mathrm{n}}(\mathbf{r};t) = 0.94e^{-\frac{\mathrm{V_m}(\mathbf{r};t)+75}{17}}$<br>$\mathrm{n}(\mathbf{r};\infty) = \frac{\alpha_{\mathrm{n}}(\mathbf{r};0)}{\alpha_{\mathrm{n}}(\mathbf{r};0)+\beta_{\mathrm{n}}(\mathbf{r};0)}$, $\tau(\mathbf{r};t) = \frac{1}{\alpha_{\mathrm{n}}(\mathbf{r};t)+\beta_{\mathrm{n}}(\mathbf{r};t)}$<br>$\alpha_{\overline{\mathrm{n}}}(\mathbf{r};t) = 0.000457e^{-\frac{\mathrm{V_m}(\mathbf{r};t)+13}{50}}$, $\beta_{\overline{n}}(\mathbf{r};t) = 0.0065e^{-\frac{\mathrm{V_m}(\mathbf{r};t)+15}{28}+1}$<br>$\overline{\mathrm{n}}(\mathbf{r};\infty) = \frac{\alpha_{\overline{\mathrm{n}}}(\mathbf{r};0)}{\alpha_{\overline{\mathrm{n}}}(\mathbf{r};0)+\beta_{\overline{n}}(\mathbf{r};0)}$, $\overline{\tau}(\mathbf{r};t) = \frac{1}{\alpha_{\overline{\mathrm{n}}}(\mathbf{r};t)+\beta_{\overline{n}}(\mathbf{r};t)}$ | $\gamma = 2, \eta = 1$ |
| $\mathrm{Ca_{LVA}}$ | $\mathrm{n}(\mathbf{r};\infty) = \frac{1}{1+e^{-\frac{\mathrm{V_m}(\mathbf{r};0)+30}{6}}}$, $\tau(\mathbf{r};t) = \frac{1}{(2.3)^{1.3}}\left[5.0 + \frac{20.0}{1+e^{\frac{\mathrm{V_m}(\mathbf{r};t)+25.0}{5.0}}}\right]$<br>$\overline{\mathrm{n}}(\mathbf{r};\infty) = \frac{1}{1+e^{\frac{\mathrm{V_m}(\mathbf{r};0)+80}{6.4}}}$, $\overline{\tau}(\mathbf{r};t) = \frac{1}{(2.3)^{1.3}}\left[20.0 + \frac{50.0}{1+e^{\frac{\mathrm{V_m}(\mathbf{r};t)+40.0}{7.0}}}\right]$ | $\gamma = 2, \eta = 1$ |
| $\mathrm{SK_{v3}}$ | $\mathrm{n}(\mathbf{r};\infty) = \frac{1}{1+e^{\frac{\mathrm{V_m}(\mathbf{r};0)-18.70}{-9.70}}}$, $\tau(\mathbf{r};t) = \frac{4.0}{1+e^{-\frac{\mathrm{V_m}(\mathbf{r};t)+46.56}{44.140}}}$ | $\gamma = 1, \eta = 0$ |
| $\mathrm{SK_{E2}}$ | $\mathrm{n}(\mathbf{r};\infty) = \frac{1}{\left(1+\frac{0.00043}{5\times10^{-5}}\right)^{4.8}}$, $\tau(\mathbf{r};t) = 1.0$ | $\gamma = 1, \eta = 0$ |
| Nap | $\alpha_{\mathrm{n}}(\mathbf{r};t) = \frac{0.182(\mathrm{V_m}(\mathbf{r};t)+38.0)}{1-e^{-\frac{\mathrm{V_m}(\mathbf{r};t)+38.0}{6.0}}}$, $\beta_{\mathrm{n}}(\mathbf{r};t) = \frac{-0.124(\mathrm{V_m}(\mathbf{r};t)+38.0)}{1-e^{\frac{\mathrm{V_m}(\mathbf{r};t)+38.0}{6.0}}}$<br>$\mathrm{n}(\mathbf{r};\infty) = \frac{1}{1+e^{-\frac{\mathrm{V_m}(\mathbf{r};0)+52.6}{4.6}}}$, $\tau(\mathbf{r};t) = \frac{6}{(2.3)^{1.3}} \cdot \frac{1}{\alpha_{\mathrm{n}}(\mathbf{r};t)+\beta_{\mathrm{n}}(\mathbf{r};t)}$<br>$\alpha_{\overline{\mathrm{n}}}(\mathbf{r};t) = \frac{-2.88\times10^{-6}(\mathrm{V_m}(\mathbf{r};t)+17.0)}{1-e^{\frac{\mathrm{V_m}(\mathbf{r};t)+17.0}{4.63}}}$<br>$\beta_{\overline{n}}(\mathbf{r};t) = \frac{6.94\times10^{-6}(\mathrm{V_m}(\mathbf{r};t)+64.4)}{1-e^{-\frac{\mathrm{V_m}(\mathbf{r};t)+64.4}{2.63}}}$<br>$\overline{\mathrm{n}}(\mathbf{r};\infty) = \frac{1}{1+e^{-\frac{\mathrm{V_m}(\mathbf{r};0)+48.8}{10}}}$, $\overline{\tau}(\mathbf{r};t) = \frac{1}{(2.3)^{1.3}} \cdot \frac{1}{\alpha_{\overline{\mathrm{n}}}(\mathbf{r};t)+\beta_{\overline{n}}(\mathbf{r};t)}$ | $\gamma = 3, \eta = 1$ |
| $\mathrm{K_{pst}}$ | $\mathrm{n}(\mathbf{r};\infty) = \frac{1}{1+e^{-\frac{\mathrm{V_m}(\mathbf{r};0)+1}{12}}}$<br>$\tau(\mathbf{r};t) = \begin{cases} \frac{1}{(2.3)^{1.3}}\left[1.25 + 175.03e^{0.026\mathrm{V_m}(\mathbf{r};t)}\right], \\ \qquad \text{if } \mathrm{V_m}(\mathbf{r};t) < -50.0\mathrm{mV} \\ \frac{1}{(2.3)^{1.3}}\left[1.25 + 13e^{-0.026\mathrm{V_m}(\mathbf{r};t)}\right], \\ \qquad \text{otherwise} \end{cases}$<br>$\overline{\mathrm{n}}(\mathbf{r};\infty) = \frac{1}{1+e^{-\frac{\mathrm{V_m}(\mathbf{r};0)+54}{-11}}}$<br>$\overline{\tau}(\mathbf{r};t) = \frac{1}{(2.3)^{1.3}}\left[360 + \left(1010 + 24(\mathrm{V_m}(\mathbf{r};t) + 55.0)\right)e^{-\left(\frac{\mathrm{V_m}(\mathbf{r};t)+75.0}{48.0}\right)^2}\right]$ | $\gamma = 2, \eta = 1$ |
| $\mathrm{K_{tst}}$ | $\mathrm{n}(\mathbf{r};\infty) = \frac{1}{1+e^{-\frac{\mathrm{V_m}(\mathbf{r};0)}{19}}}$<br>$\tau(\mathbf{r};t) = \frac{1}{(2.3)^{1.3}}\left[0.34 + 0.92e^{-\left(\frac{\mathrm{V_m}(\mathbf{r};t)+71.0}{59.0}\right)^2}\right]$<br>$\overline{\mathrm{n}}(\mathbf{r};\infty) = \frac{1}{1+e^{\frac{\mathrm{V_m}(\mathbf{r};0)+66}{10}}}$<br>$\overline{\tau}(\mathbf{r};t) = \frac{1}{(2.3)^{1.3}}\left[8.0 + 49.0e^{-\left(\frac{\mathrm{V_m}(\mathbf{r};t)+73.0}{23.0}\right)^2}\right]$ | $\gamma = 4, \eta = 1$ |
| NaTa | $\alpha_{\mathrm{n}}(\mathbf{r};t) = \frac{0.182(\mathrm{V_m}(\mathbf{r};t)+38)}{1-e^{-\frac{\mathrm{V_m}(\mathbf{r};t)+38}{6}}}$, $\beta_{\mathrm{n}}(\mathbf{r};t) = \frac{-0.124(\mathrm{V_m}(\mathbf{r};t)+38)}{1-e^{\frac{\mathrm{V_m}(\mathbf{r};t)+38}{6}}}$<br>$\mathrm{n}(\mathbf{r};\infty) = \frac{\alpha_{\mathrm{n}}(\mathbf{r};0)}{\alpha_{\mathrm{n}}(\mathbf{r};0)+\beta_{\mathrm{n}}(\mathbf{r};0)}$, $\tau(\mathbf{r};t) = \frac{1}{(2.3)^{1.3}} \cdot \frac{1}{\alpha_{\mathrm{n}}(\mathbf{r};t)+\beta_{\mathrm{n}}(\mathbf{r};t)}$<br>$\alpha_{\overline{\mathrm{n}}}(\mathbf{r};t) = \frac{-0.015(\mathrm{V_m}(\mathbf{r};t)+66)}{1-e^{\frac{\mathrm{V_m}(\mathbf{r};t)+66}{6}}}$, $\beta_{\overline{n}}(\mathbf{r};t) = \frac{0.015(\mathrm{V_m}(\mathbf{r};t)+66)}{1-e^{-\frac{\mathrm{V_m}(\mathbf{r};t)+66}{6}}}$<br>$\overline{\mathrm{n}}(\mathbf{r};\infty) = \frac{\alpha_{\overline{\mathrm{n}}}(\mathbf{r};0)}{\alpha_{\overline{\mathrm{n}}}(\mathbf{r};0)+\beta_{\overline{n}}(\mathbf{r};0)}$, $\overline{\tau}(\mathbf{r};t) = \frac{1}{(2.3)^{1.3}} \cdot \frac{1}{\alpha_{\overline{\mathrm{n}}}(\mathbf{r};t)+\beta_{\overline{n}}(\mathbf{r};t)}$ | $\gamma = 3, \eta = 1$ |
| Ih | $\alpha_{\mathrm{n}}(\mathbf{r};t) = \frac{0.00643(\mathrm{V_m}(\mathbf{r};t)+154.9)}{e^{\frac{\mathrm{V_m}(\mathbf{r};t)+154.9}{11.9}}-1}$, $\beta_{\mathrm{n}}(\mathbf{r};t) = 0.193e^{\frac{\mathrm{V_m}}{33.1}}$<br>$\mathrm{n}(\mathbf{r};\infty) = \frac{\alpha_{n}(\mathbf{r};0)}{\alpha_{n}(\mathbf{r};0)+\beta_{n}(\mathbf{r};0)}$, $\tau_{\mathrm{n}}(\mathbf{r};t) = \frac{1}{\alpha_{\mathrm{n}}(\mathbf{r};t)+\beta_{\mathrm{n}}(\mathbf{r};t)}$ | $\gamma = 1, \eta = 0$ |
| Im | $\alpha_{\mathrm{n}}(\mathbf{r};t) = 0.0033e^{0.1(\mathrm{V_m}(\mathbf{r};t)+35.0)}$<br>$\beta_{\mathrm{n}}(\mathbf{r};t) = 0.0033e^{-0.1(\mathrm{V_m}(\mathbf{r};t)+35.0)}$<br>$\mathrm{n}(\mathbf{r};\infty) = \frac{\alpha_{n}(\mathbf{r};0)}{\alpha_{n}(\mathbf{r};0)+\beta_{n}(\mathbf{r};0)}$ | $\gamma = 1, \eta = 0$ |
| NaTs2 | $\alpha_{\mathrm{n}}(\mathbf{r};t) = \frac{0.182(\mathrm{V_m}(\mathbf{r};t)+32.0)}{1-e^{-\frac{\mathrm{V_m}(\mathbf{r};t)+32.0}{6.0}}}$, $\beta_{\mathrm{n}}(\mathbf{r};t) = \frac{-0.124(\mathrm{V_m}(\mathbf{r};t)+32.0)}{1-e^{-\frac{\mathrm{V_m}(\mathbf{r};t)+32.0}{6.0}}}$<br>$\mathrm{n}(\mathbf{r};\infty) = \frac{\alpha_{\mathrm{n}}(\mathbf{r};0)}{\alpha_{\mathrm{n}}(\mathbf{r};0)+\beta_{\mathrm{n}}(\mathbf{r};0)}$, $\tau_{\mathrm{n}}(\mathbf{r};t) = \frac{1}{(2.3)^{1.3}} \cdot \frac{1}{\alpha_{\mathrm{n}}(\mathbf{r};t)+\beta_{\mathrm{n}}(\mathbf{r};t)}$<br>$\alpha_{\overline{\mathrm{n}}}(\mathbf{r};t) = \frac{-0.015(\mathrm{V_m}(\mathbf{r};t)+60.0)}{1-e^{\frac{\mathrm{V_m}(\mathbf{r};t)+60.0}{6.0}}}$, $\beta_{\overline{n}}(\mathbf{r};t) = \frac{0.015(\mathrm{V_m}(\mathbf{r};t)+60.0)}{1-e^{-\frac{\mathrm{V_m}(\mathbf{r};t)+60.0}{6.0}}}$<br>$\overline{\mathrm{n}}(\mathbf{r};\infty) = \frac{\alpha_{\overline{\mathrm{n}}}(\mathbf{r};0)}{\alpha_{\overline{\mathrm{n}}}(\mathbf{r};0)+\beta_{\overline{n}}(\mathbf{r};0)}$, $\tau_{\overline{n}}(\mathbf{r};t) = \frac{1}{(2.3)^{1.3}} \cdot \frac{1}{\alpha_{\overline{\mathrm{n}}}(\mathbf{r};t)+\beta_{\overline{n}}(\mathbf{r};t)}$ | $\gamma = 3, \eta = 1$ |

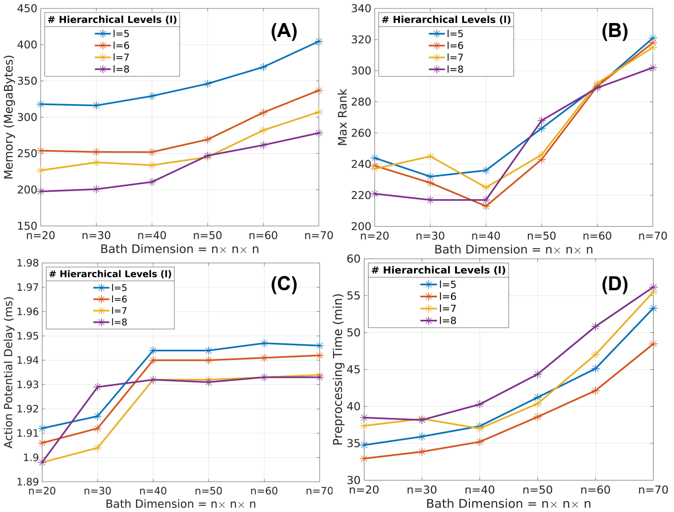


Fig. 17: Effects of the bath size ($n \times n \times n$) and the number of HODLR hierarchical levels ($l$) on the computational memory of HODLR compression (A), the HODLR maximum rank (B), initialization of the action potential (C), and total compression time (D).

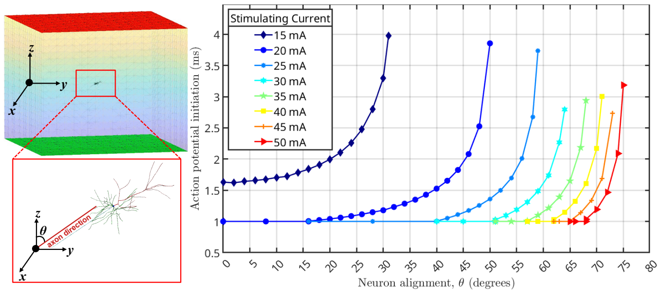


Fig. 18: Effect of neuron spatial orientation on action potential initiation. For the same alignment, the lower stimulating current results in a slower initiation than that of the higher stimulation current.

#### S8. Effect of nearby neurons on action potential initiation

Fig. [20] shows the effect of a nearby second neuron ($N_2$) on the initiation of the action potential in the observed neuron ($N_1$). The second neuron has a negligible effect on the observed neuron.

#### S9. Effect of ion mechanism on the action potential

Fig. [21] shows how the action potential changes across different ion mechanisms. For this purpose, the action potential is sampled along the axon (sample points $P_1$-$P_7$) and the soma (sample point $P_8$) as shown in Fig. [21]. Fig. [21B] shows that the membrane voltage ($V_m$) propagates without relaxing. As the action potential propagates into the soma, there's a significant change in the ionic currents ($I_{ion}$), total ionic conductance ($\sum_i g_i$), charge density ($\rho$) and the membrane current ($I_m$); thereby, affecting the membrane voltage.

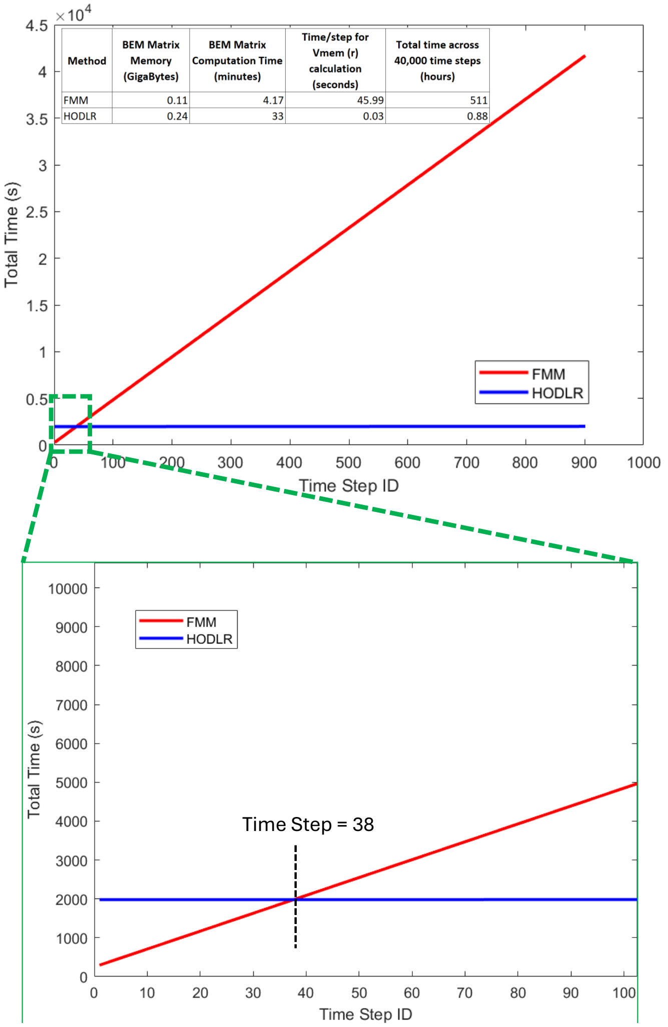


Fig. 19: The FMM iterative solver has lower computational time for the first few time-steps than the HODLR compression, but scales linearly across 40,000 time steps, therefore, making HODLR compression a better choice than FMM.

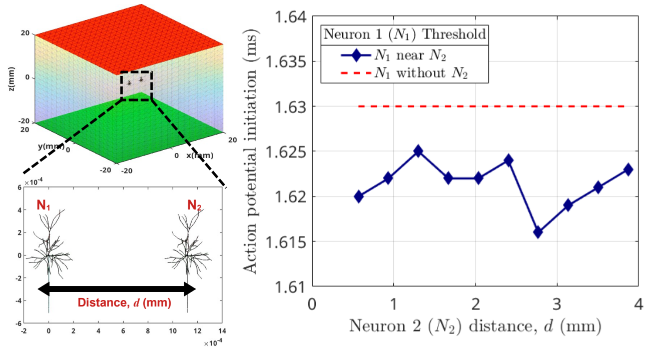


Fig. 20: Effect of nearby neurons on action potential initiation. The second neuron ($N_2$) has a negligible effect on the observed neuron ($N_1$).

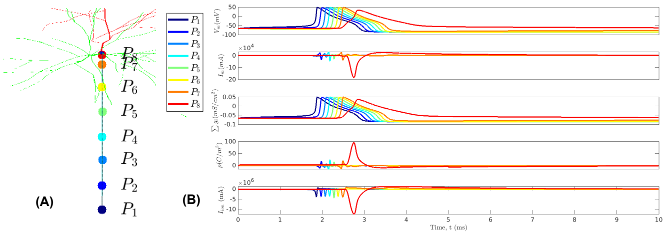

Fig. 21: Effect of ion mechanism on the propagation of action potential. The membrane voltage propagates without any relaxation through the axon (sampled points $P_1$-$P_7$). Inside the soma, the action potential changes due to different ion mechanisms. Similar effects are shown in the ionic currents ($I_{ion}$), total ionic conductance ($\sum_i g_i$), charge density ($\rho$) and the membrane current ($I_m$).
